# A bipartite transcription factor module controlling expression in the bundle sheath of *Arabidopsis thaliana*

**DOI:** 10.1101/380188

**Authors:** Patrick J. Dickinson, Jana Kneřová, Marek Szecówka, Sean S. Stevenson, Steven J. Burgess, Hugh Mulvey, Anne-Maarit Bågman, Allison Gaudinier, Siobhan M. Brady, Julian M. Hibberd

## Abstract

C_4_ photosynthesis evolved repeatedly from the ancestral C_3_ state, improving photosynthetic efficiency by ∼50%. In most C_4_ lineages photosynthesis is compartmented between mesophyll and bundle sheath cells but how gene expression is restricted to these cell types is poorly understood. Using the C_3_ model *Arabidopsis thaliana* we identified *cis*-elements and transcription factors driving expression in bundle sheath strands. Upstream of the bundle sheath preferentially expressed *MYB76* gene we identified a region necessary and sufficient for expression containing two *cis*-elements associated with the MYC and MYB families of transcription factors. *MYB76* expression is reduced in mutant alleles for each. Moreover, down-regulated genes shared by both mutants are preferentially expressed in the bundle sheath. Our findings are broadly relevant for understanding the spatial patterning of gene expression, provide specific insights into mechanisms associated with evolution of C_4_ photosynthesis and identify a short tuneable sequence for manipulating gene expression in the bundle sheath.

## Introduction

A fundamental characteristic of multicellular eukaryotes is the ability to carry out diverse and specialised functions in distinct tissues. Diversification in tissue function is associated with variation in protein content that is to a large extent determined by patterns of gene expression. One striking example of metabolic compartmentalisation is represented by C_4_ photosynthesis where carbon is initially fixed in the mesophyll but is then released and re-fixed in bundle sheath cells. The C_4_ pathway is more efficient than C_3_ photosynthesis under warm, dry conditions, and as a consequence it has been proposed that engineering the C_4_ pathway into C_3_ crops such as rice could lead to increased yields (Hibberd et al., 2008; Von Caemmerer et al., 2012). Understanding mechanisms directing expression to bundle sheath or mesophyll cells is crucial to this effort and previous work has shown that multiple mechanisms can drive cell type-preferential expression in C_4_ species (Hibberd & Covshoff, 2010). For example, expression of the *Glycine decarboxylase P subunit (GLDPA)* in the bundle sheath and veins of C_4_ *Flaveria bidentis* is due to interplay between multiple regulatory regions (Wiludda et al., 2012). Sequence in the distal promoter generates strong expression but is not tissue-specific, however in the presence of proximal promoter elements, expression in the bundle sheath is brought about by transcripts derived from the distal promoter being degraded in mesophyll cells through nonsense-mediated RNA decay of incompletely spliced transcripts (Engelmann et al., 2008; Wiludda et al., 2012). Similarly, for the *Phosphoenolpyruvate carboxylaseA1 (PpcA1)* gene from C_4_ *Flaveria trinervia*, two submodules in a distal region that are enhanced by interaction with sequence in the proximal promoter are sufficient to confer mesophyll specificity (Gowik et al., 2004; Akyildiz et al., 2007). In addition to promoter sequences, other genic regions contain *cis-*elements that generate tissue-specific gene expression. For example, preferential expression of the *CARBONIC ANHYDRASE2, CARBONIC ANHYDRASE4* and *PYRUVATE,ORTHOPHOSPHATE DIKINASE* genes in mesophyll cells of the C_4_ species *Gynandropsis gynandra* is mediated by a nine base pair motif present in both 5’ and 3’ untranslated regions (Williams et al., 2016). Moreover, preferential expression of *NAD-ME1&2* genes in the bundle sheath of *G. gynandra* is associated with two motifs known as Bundle Sheath Modules (BSM) 1a and 1b that co-operatively restrict gene expression to this tissue. BSM1a and BSM1b represent duons because they are located in coding sequence and so determine amino acid composition as well as gene expression (Brown et al., 2011; Reyna-Llorens et al., 2018). In summary, tissue-specific expression can be generated through multiple mechanisms, but factors in *trans* that interact with the *cis*-elements controlling tissue specific patterning of gene expression have not yet been identified.

As the C_4_ pathway appears to have evolved repeatedly from the ancestral C_3_ state by co-opting existing molecular mechanisms from C_3_ leaves (Matsuoka et al., 1994: Kajala et al., 2012; Reyna-Llorens & Hibberd, 2017) we sought to leverage the C_3_ model *Arabidopsis thaliana* (hereafter Arabidopsis) to better understand mechanisms allowing cell type-specific gene expression. The bundle sheath represents about 15% of cells in leaves of Arabidopsis (Kinsman & Pyke, 1998) and has been proposed to play important roles in hydraulic conductance (Shatil-Cohen et al., 2011), transport of metabolites (Leegood, 2008), as well as storage of carbohydrates (Koroleva et al., 1997), ions (Williams et al., 2018) and water (Sage, 2001; Griffiths et al., 2013). A number of findings are consistent with bundle sheath cells also being involved in sulphur metabolism and glucosinolate biosynthesis. First, the promoter of *SULPHUR TRANSPORTER2.2* generates preferential expression in the bundle sheath (Takahashi et al., 2000; Kirschner et al., 2018) and secondly, compared with the whole leaf, transcripts encoding enzymes of sulphur metabolism are more abundant on bundle sheath ribosomes (Aubry et al., 2014). Transcripts of *MYB76* and other MYB domain transcription factors involved in glucosinolate biosynthesis are also preferentially associated with bundle sheath ribosomes (Aubry et al., 2014) but how this patterning of gene expression is achieved is not known.

To better understand the regulation of cell type-specific gene expression in Arabidopsis we focussed on the *MYB76* gene that is preferentially expressed in the bundle sheath. A classical truncation analysis was combined with computational interrogation of transcription factor binding sites to identify a 256-nucleotide region necessary and sufficient for expression in the Arabidopsis bundle sheath. Within this region we identified MYC and MYB transcription factor binding sites. We show MYC and MYB transcription factors are necessary for *MYB76* expression as well as the expression of at least forty-seven additional genes that are preferentially expressed in the bundle sheath. We propose that the MYC-MYB module previously associated with expression of glucosinolate biosynthetic genes (Schweizer et al., 2013) acts as a driver of bundle sheath preferential expression in Arabidopsis. To our knowledge, this work provides the first example of a regulatory system governing the spatial control of gene expression in leaves.

## Results

### A short region in the *MYB76* promoter necessary for expression in the bundle sheath

Gene Ontology (GO) analysis of publicly available data (Aubry et al., 2014) indicated that in addition to transcripts encoding proteins important for amino acid export, those encoding glucosinolate (GLS) biosynthesis proteins are strongly enriched in the bundle sheath of Arabidopsis (Figure 1A). In fact, all but five of the thirty genes reported to be involved in GLS biosynthesis (Li et al., 2014) showed more than two-fold higher expression in the bundle sheath compared with the whole leaf (Figure 1B). The expression of genes involved in aliphatic GLS metabolism is mostly controlled by MYC and MYB transcription factors (Gigolashvili et al., 2008; Schweizer et al., 2013) including MYC2, MYC3, MYC4 and MYB28, MYB29 and MYB76. Notably, genes encoding MYB28, MYB29 and MYB76 were strongly expressed in the bundle sheath compared with the whole leaf (Figure 1B). We sought to use these transcription factors to better understand how gene expression is restricted to the bundle sheath of C_3_ plants.

**Figure 1:**
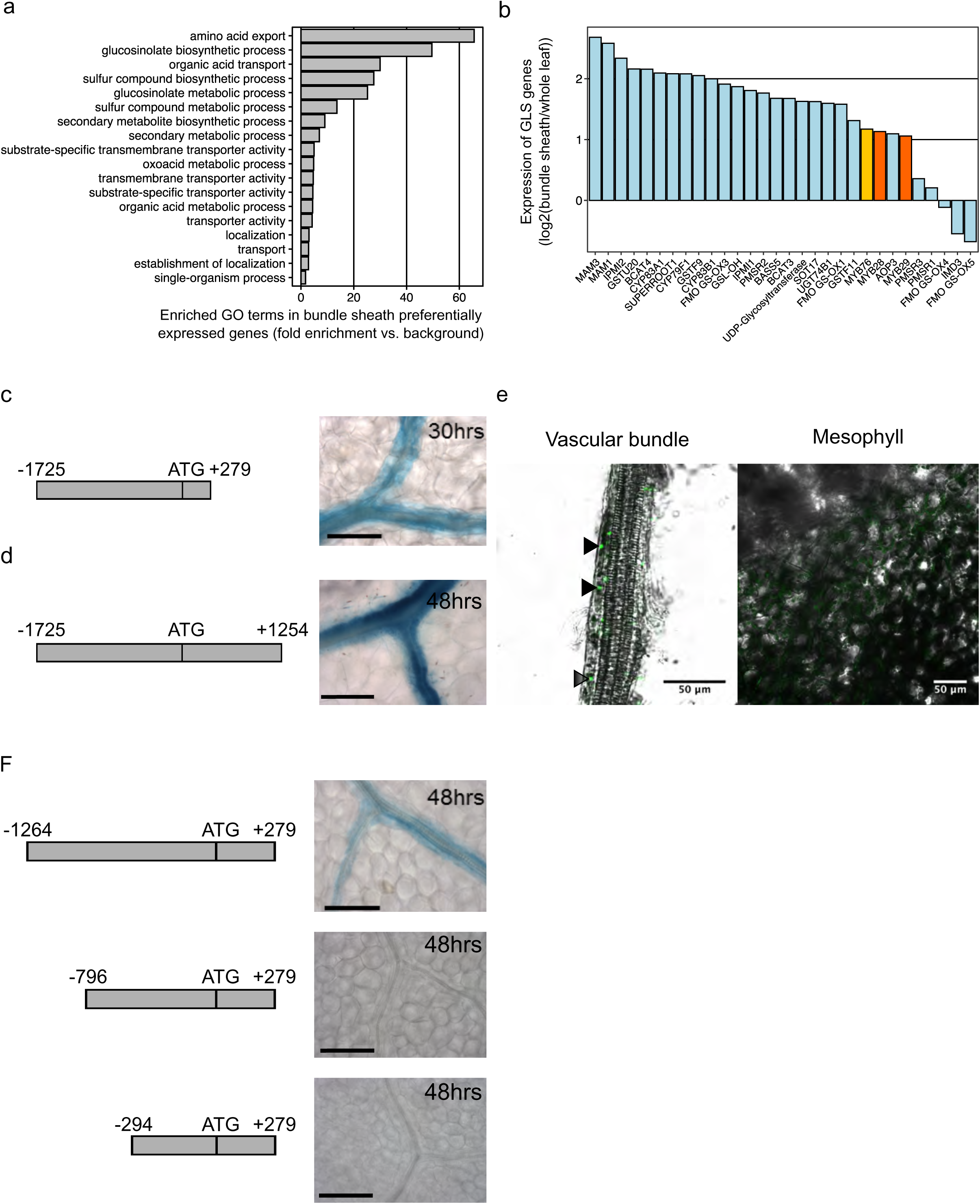
A 468bp region from the *MYB76* promoter necessary for bundle-sheath expression. A) Gene Ontology term enrichment of the 200 most bundle sheath preferential genes in Arabidopsis. Enrichment shown as fold enrichment compared with background. B) Expression of glucosinolate biosynthesis genes in bundle sheath compared with whole leaves. *MYB76* marked in gold, *MYB28* and *MYB29* marked in orange. C) Schematic and representative image of the *MYB76* promoter plus 279bp of genomic sequence fused to GUS. D) Schematic and representative image of the *MYB76* promoter and full genomic sequence fused to GUS. For C and D staining times are given in the top right corner of each image and scale bars represent 100μm. E) Representative images of the *MYB76* promoter driving expression of a histone GFP fusion. Images of the vascular bundle (left) and mesophyll (right). Black arrowheads indicate nuclei expressing GFP. F) Schematics and representative GUS images of each *MYB76* deletion. Staining times are given in the top right corner of each image. Scale bars represent 100μm.

To test whether regions upstream of *MYB28, MYB29* and *MYB76* were sufficient to drive expression in the bundle sheath of Arabidopsis, each was fused to the *uidA* reporter gene encoding GUS and multiple transgenic lines were generated. We were unable to detect GUS staining in leaves from the promoter of *MYB28* alone (Supplemental Figure 1) and whilst the promoter of *MYB29* did mark veins and bundle sheath cells, it also led to some GUS accumulating in mesophyll cells (Supplemental Figure 1). In contrast, a construct containing the promoter and 279bp after the translational start site of *MYB76* generated clearly detectable GUS in the Arabidopsis bundle sheath with no GUS detected in the mesophyll (Figure 1C, Supplemental Figure 2). We conclude that regulatory elements sufficient for bundle sheath expression of *MYB76* (Figure 1B) are contained in this sequence but the preferential accumulation of transcripts from *MYB28* and *MYB29* (Supplemental Figure 1) is likely mediated by *cis*-elements located outside of the sequence tested.

As expression patterns can be determined by *cis*-elements in promoters, untranslated regions, exons, introns or downstream 3’ regions (Ali & Taylor, 2001; Brown et al., 2011; Kajala et al., 2011; Williams et al., 2016; Gallegos & Rose, 2017) a translational fusion between the *MYB76* genomic sequence and *uidA* driven by the *MYB76* promoter was generated to confirm that the strong expression in the bundle sheath mediated by the *MYB76* promoter reflected the pattern of expression associated with the intact genomic sequence. Transgenic lines harbouring this genomic fusion showed preferential accumulation of GUS in the bundle sheath (Figure 1D, Supplemental Figure 3) mirroring the pattern found from the promoter alone. Use of the fluorometric 4-MethylUmbelliferyl β-D-Glucuronide (MUG) assay showed that GUS accumulation was lower when nucleotides +280 to +1254 relative to the translational start site were included (Supplemental Figure 6) suggesting that the full genomic sequence of *MYB76* contains regulators that quantitatively repress expression. To confirm that the GUS reporter generated a reliable read-out of spatial expression patterns, a nuclear localised *pMYB76::H2B::GFP* line was produced. Imaging of GFP in deep tissue of leaves such as the bundle sheath is challenging, but consistent with data from the GUS reporter (Figure 1C), GFP was detectable in nuclei of the vasculature and bundle sheath cells but was absent from the mesophyll (Figure 1E, Supplemental Figure 4). These results show that the promoter of *MYB76* generates bundle sheath preferential expression.

To further investigate the elements driving *MYB76* expression in the bundle sheath, 5’ deletions of the promoter were generated. Removal of nucleotides -1725 to -1264 relative to the translational start site did not impact on GUS localisation, however once nucleotides -1264 to -796 were removed GUS was no longer detectable in the bundle sheath. Further removal of another 500 base pairs had no additional impact on the spatial pattern of GUS accumulation (Fig 1F, Supplemental Figure 5). These findings are supported by quantification *via* MUG assays (Supplemental Figure 6) that showed removal of nucleotides -1725 to -1264 reduced accumulation of the reporter, and MUG was no longer detectable once sequence upstream of nucleotide -796 was absent. Overall, these data indicate that the *MYB76* promoter contains a region between nucleotides -1264 and -796 upstream of the translational start site that directs expression to the bundle sheath.

### A DNaseI Hypersensitive Site in the *MYB76* promoter is necessary and sufficient for bundle sheath preferential expression

The DNaseI enzyme preferentially cuts accessible DNA and so can be used to define sequences available for transcription factor binding and thus the location of regulatory DNA (Hesselberth et al., 2009). To complement our truncation analysis, an existing dataset that defined DNaseI Hypersensitive Sites (DHS) in Arabidopsis (Zhang et al., 2012) was interrogated. Two DHS were detected upstream of *MYB76* in both flower tissue and leaves (Figure 2A) with a DHS encompassing nucleotides -909 to -654 upstream of the translational start site overlapping with the region required for expression in the bundle sheath (Figure 1F). Consistent with the DHS data, *MYB76* has been reported to be expressed in both leaves and flowers (Gigolashvili et al. 2008). Although the DHS had a lower DHS score in the leaf than the flower (mean DHS score of 1.3 versus 2.6 (Zhang et al., 2016)), this may be due to the fact that bundle sheath cells make up a small proportion of cells in a C_3_ leaf (Kinsman & Pyke, 1998) such that transcription factor binding upstream of *MYB76* would be diluted in whole leaves.

**Figure 2:**
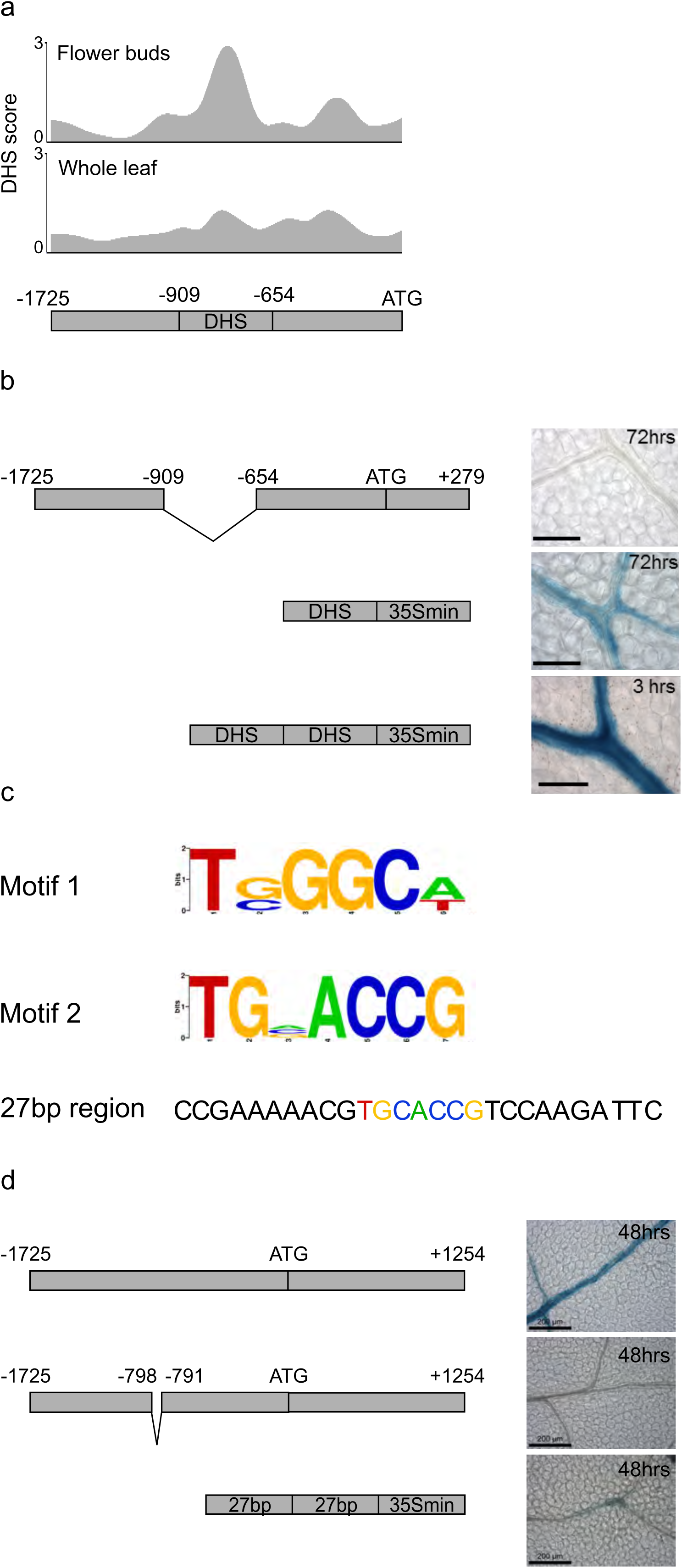
A DNaseI Hypersensitive Site in the *MYB76* promoter is necessary and sufficient for expression in the bundle-sheath. A) The *MYB76* promoter contains a DNaseI Hypersensitive Site (DHS) located between nucleotides -909 to -654. The y-axis shows the DHS score (Zhang et al., 2016) from flower buds (top) and leaves (bottom). Data are from Zhang et al., (2012) visualised with the IGV browser (Robinson et al., 2011). B) Schematics and representative GUS staining images of the *MYB76* promoter with the DHS deleted, the DHS fused to the minimal *CaMV35S* promoter and two copies of the DHS fused to the minimal *CaMV35S* promoter. Staining times are given in the top right corner of each image. Scale bars represent 100μm. C) Position Weight Matrices (PWMs) of two motifs (1 and 2) found in the *MYB76* DHS as well as other promoters driving bundle sheath expression in Arabidopsis and a 27bp region of the *MYB76* DHS containing motif 2 (TGCACCG and highlighted in colours matching the PWM). D) Schematics and representative images of *MYB76* promoter and gDNA (top), *MYB76* promoter and genomic DNA with the *TGCACCG* motif deleted (middle), and oligomerisation of the 27bp region containing TGCACCG (bottom) fused to GUS. Staining times are given in the top right corner of each image. Scale bars represent 200μm.

Removing the DHS found between nucleotides -909 to -654 of the *MYB76* promoter abolished accumulation of GUS, and fusing this DHS to the minimal *CaMV35S* promoter was sufficient to generate GUS in the bundle sheath (Figure 2B, Supplemental Figure 7). Furthermore, oligomerizing two copies of the DHS upstream of the minimal *CaMV35S* promoter resulted in very strong accumulation of GUS in the bundle sheath (Figure 2B, Supplemental Figure 7). From these data we conclude that sequence within this DHS is both necessary and sufficient to activate expression preferentially in the bundle sheath. Combined with the truncation analysis indicating that nucleotides -1264 to -796 upstream of the translational start site were required for expression in the bundle sheath (Figure 1F, Supplemental Figure 5), our findings indicate that a positive regulator of bundle sheath expression is located between nucleotides -909 (the start of the DHS) and -796 upstream of the *MYB76* translational start site.

Phylogenetic foot-printing identified two motifs (Figure 2C) in the *MYB76* DHS that are shared by *MYB76* and the promoters of *SCR, SULTR2.2* and *GLDP* that have previously been reported to generate expression in the Arabidopsis bundle sheath (Wysocka-Diller et al. 2000; Engelmann et al. 2008; Takahashi et al. 2000; Kirschner et al. 2018). Whilst site directed mutagenesis of motif one (TGGGCA) had no impact on accumulation of GUS in the bundle sheath (Supplemental Figure 8) deletion of motif two (TGCACCG) in the context of the full genomic sequence of *MYB76* abolished GUS accumulation in the bundle sheath (Figure 2D, Supplemental Figure 8). These data indicate that this sequence is necessary to pattern expression from both the promoter of *MYB76* alone, but also the full genic *MYB76* sequence containing exons, introns and UTRs. To test whether this sequence is sufficient to direct expression to the bundle sheath it was combined with ten upstream and ten downstream nucleotides from the endogenous *MYB76* promoter, oligomerized, and fused to *uidA*. Although this construct did not recapitulate the strong bundle sheath expression of the *MYB76* DHS it was able to generate weak expression in the bundle sheath (Figure 2D, Supplemental Figure 10). We conclude that the 27bp sequence is necessary and weakly sufficient to direct expression to the bundle sheath.

### MYC, MYB and DREB transcription factors control bundle sheath expression of *MYB76* from the DHS

To better understand the *cis*-regulatory landscape within the *MYB76* DHS we used the Find Individual Motif Occurrences (FIMO) tool (Grant et al., 2011) to predict transcription factor binding sites. For the majority of Arabidopsis transcription factors, DNA binding sites have not yet been defined, and so to allow us to search for broad consensus sequences associated with groups of transcription factors we clustered the 555 transcription factor binding motifs for Arabidopsis from the JASPAR motif database (Formes et al., 2019) into 43 groups based on relatedness of the motif position weight matrices (PWMs) (Supplemental Table 1). Plotting matches for each of these motifs in the *MYB76* DHS (Figure 3A) showed that it contained binding sites for motifs from twelve clusters. Notably the 27bp region necessary for expression in the bundle sheath contained binding sites from clusters 1, 8, 11 and 16 that correspond to the binding sites from ERF (clusters 1 and 11), bHLH (cluster 8), and G2-like (cluster 16) transcription factor families. There are also predicted binding sites for IDD (cluster 9), MADS (cluster 23) and MYB (clusters 10 and 18) families within the DHS. To supplement this *in silico* analysis, Yeast One-Hybrid identified thirteen transcription factors from seven different families that were able to bind the DHS (Figure 3B). Six of these transcription factors including DF1, MYB73 and AIL5 were previously reported to bind the whole *MYB76* promoter (Li et al. 2014) (Figure 3B). Although not identified by Yeast One-Hybrid, MYC2, MYC3, MYC4, MYB28 and MYB29 can control *MYB76* expression (Gigolashvili et al., 2008; Schweizer et al., 2013) and so were incorporated into our list of candidate regulators of *MYB76*. Of these candidates, *MYB28*, *MYB29*, *DF1*, *MYB73* and *AIL5* were strongly preferentially expressed in the bundle sheath (with a log2(bundle sheath/whole leaf) > 0.75 cut off) (Figure 3B).

**Figure 3:**
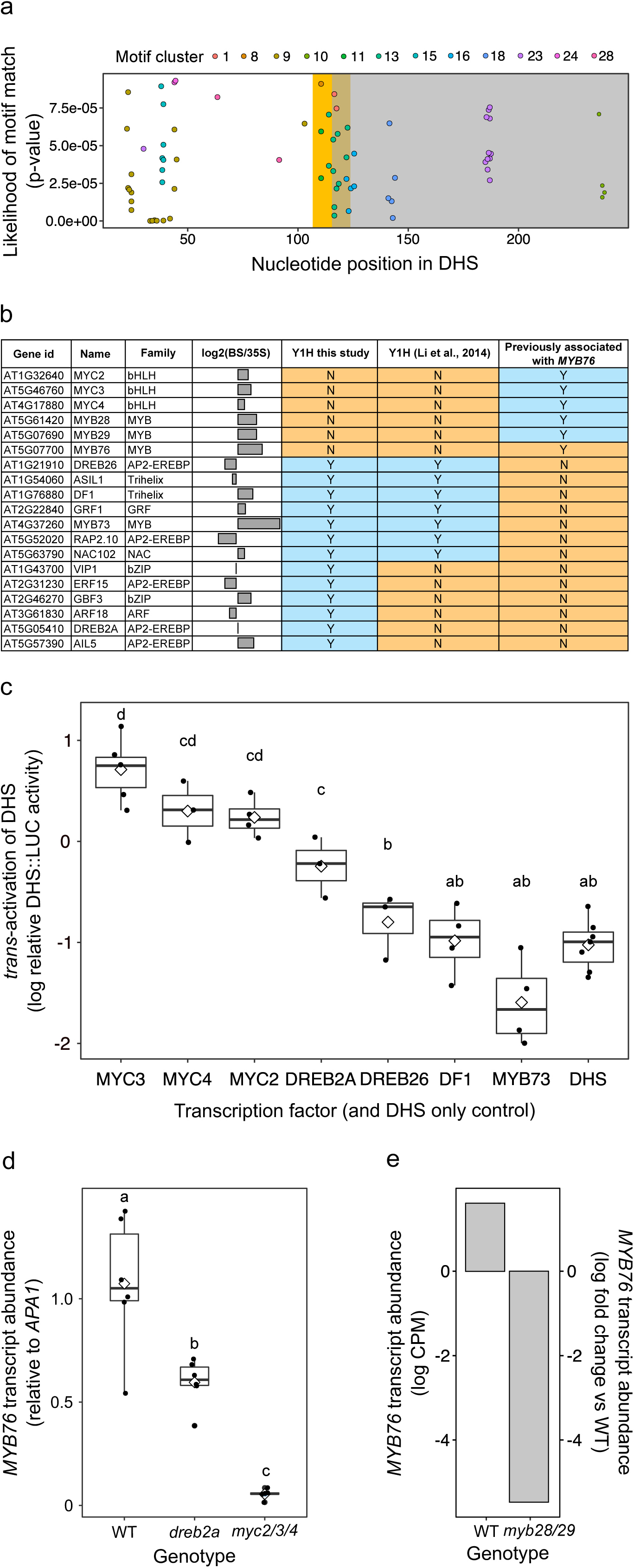
MYC, MYB and DREB transcription factors control *MYB76* expression from the DHS. A) Transcription factor binding motifs within the *MYB76* DHS. Position in the DHS (bp) is on the x-axis, and predicted binding affinity (p-values) on the y-axis. Motifs are coloured by motif cluster (Supplemental table 1). The gold region represents the 27bp region necessary for expression and the grey region indicates sequence unable to generate bundle sheath expression (Figure 1F). B) Summary of candidate transcription factors binding to the *MYB76* DHS. Information provided includes gene identifier, gene name, family, expression in bundle sheath compared with whole leaves (from Aubry et al., 2014), whether they interacted with the DHS in Yeast One-Hybrid, whether they were previously identified as binding the entire *MYB76* promoter (from Li et al., 2014), and if they have previously been associated with controlling *MYB76* expression (Gigolashvili et al., 2008; Schweizer et al., 2013). C) *Trans-*activation assays of candidate transcription factors and the *MYB76* DHS. Values shown represent the log of LUCIFERASE (LUC) signal driven by *MYB76DHS::LUC* normalised to a constitutively expressed GUS infiltration control. Box-plots show inter-quartile range as upper and lower confines of the box, median as a solid black line, mean as a white diamond and whiskers as maximum and minimum values excluding outliers. All individual data points are plotted. a, b, c and d represent statistically significant differences (p<0.05) as determined by pairwise T-tests. n = independent biological samples with n=3 for DREB2A, DREB26 and MYC4, n=4 for MYC2, DF1 and MYB73, n=6 for MYC3 and n=7 for DHS. D) qRT-PCR of *MYB76* in WT, *dreb2a* and *myc2/3/4*. Expression shown relative to *APA1,* n=6 independent biological samples for all genotypes. Box-plots show inter-quartile range as upper and lower confines of the box, median as a solid black line, mean as a white diamond and whiskers as maximum and minimum values excluding outliers. All individual data points are plotted. All individual data points are plotted. a, b and c represent statistically significantly differences (p<0.001) determined by pairwise T-tests. E) *MYB76* expression from a publically available *myb28/29* transcriptome (Burow et al., 2015). Expression in WT (left) shown as log counts per million. Expression in *myb28/29* (right) shown as log fold change relative to wild type.

We next tested whether these candidate transcription factors could activate expression from the *MYB76* DHS *in planta.* For the following reasons we chose to test MYC2, MYC3, MYC4, DREB2A, DREB26, DF1 and MYB73. First, MYC2, MYC3, MYC4, DREB2A and DREB26 have binding sites for the clusters described above within the 27bp region and MYC2, MYC3 and MYC4 have previously been reported to affect *MYB76* expression (Schweizer et al., 2013). Second, DREB2A was found in our Yeast One-Hybrid experiment and DREB26 was identified in both Yeast One-Hybrid screens (Figure 3B). Third, DF1 and MYB73 were found in both Yeast One-Hybrid studies (Figure 3B), are preferentially expressed in the bundle sheath (Figure 3B) and DF1 has previously been associated with *GLS* gene expression (Li et al., 2014). Although the DHS contains two additional predicted MYB binding sites (one associated with cluster 10 and one with cluster 18 MYBs) these were outside the region necessary for activating *MYB76* expression in the bundle sheath (Figure 3A). Each transcription factor was used in a *trans-*activation assay for the *MYB76* DHS in *Nicotiana benthamiana.* Infiltration with MYC2, MYC3, MYC4 and DREB2A resulted in significantly more LUCIFERASE (LUC) signal than infiltration with the DHS alone, with MYC2, MYC3 and MYC4 driving higher LUC signal than DREB2A (Figure 3C). LUC signal was not significantly different from the DHS alone for any of the other transcription factors tested (Figure 3C).

As the MYCs (Cluster 8) and DREB2A (Cluster 11) have predicted binding sites that overlap the 27bp region necessary for expression in the bundle sheath (Figure 3A) and were able to activate expression from the DHS in the *trans-*activation assays (Figure 3C) we tested whether expression of *MYB76* was perturbed in mutant alleles of each. qRT-PCR on *MYB76* was performed on *myc2/3/4* and *dreb2a* mutants. *MYB76* expression was reduced by approximately half in the *dreb2a* mutant, and in the *myc2/3/4* triple mutant by about 19 times (Figure 3D). These data are consistent with the *trans-*activation results which showed a strong increase in expression driven by MYC transcription factors and a weaker increase driven by DREB2A (Figure 3C). Taken together this indicates that under the conditions we used MYC2, MYC3 and MYC4 have a major role in controlling *MYB76* expression and DREB2A has a smaller effect. Previous work has shown that MYC transcription factors interact with MYB28, MYB29 and MYB76 to activate the expression of genes involved in GLS metabolism (Schweizer et al., 2013). Therefore, despite them not appearing in either Yeast One-Hybrid screen, we asked whether they are involved in controlling *MYB76* expression. Re-analysis of publicly available data (Burow et al., 2015) showed that *MYB76* expression was substantially reduced in a *myb28/29* mutant (Figure 3E). This is consistent with previous reports of MYB28, 29 and 76 being able to activate each other’s expression (Gigolashvili et al., 2008). *MYB76* is expressed at similar levels to wild type in both *myb28* and *myb29* single mutants (Sønderby et al., 2007) suggesting that there is redundancy between *MYB28* and *MYB29* in the control of *MYB76* expression. Although *MYB29* and *MYB76* are tandem duplicates on chromosome five, because *MYB76* is expressed similarly to wild type in a *myb29* mutant, the reduction of *MYB76* expression in the *myb28/29* mutant is unlikely to be a result of the proximity of the *myb29-1* T-DNA insertion to *MYB76.* MYB28, 29 and 76 do not have defined transcription factor binding motifs in publicly available databases, however mapping motif clusters to a phylogenetic reconstruction of MYB transcription factors (Supplemental Figure 11) showed that MYB28, MYB29 and MYB76 were found in the cluster 18 clade, strongly suggesting that their binding preference is similar to those of cluster 18 MYBs (Supplemental Table 1). Taken together these data suggest that MYC2/MYC3/MYC4 as well as MYB28/MYB29 are involved in controlling *MYB76* expression from the DHS.

### MYC2/3/4 and MYB28/29 control the expression of many other bundle sheath preferential genes in Arabidopsis

The findings above are consistent with the MYC-MYB module, previously reported to activate GLS metabolism genes in response to herbivory (Schweizer et al., 2013) activating *MYB76* in the bundle sheath in the absence of herbivory. As this module is involved in activating the expression of multiple GLS metabolism genes in Arabidopsis (Schweizer et al., 2013) and most are preferentially expressed in the bundle sheath (Figure 1B), we wished to test whether the MYC-MYB system might also be responsible for their bundle sheath preferential expression. We therefore re-analysed publicly available transcriptome data for *myc2/3/4* (Major et al., 2017) and *myb28/29* (Burow et al., 2015) mutants. We identified 207 genes that were down-regulated (log2 vs. WT < -0.75) in *myc2/3/4* (Figure 4A), 729 genes that were down-regulated in *myb28/29* (Figure 4B) and 76 genes that were down-regulated in both *myc2/3/4* and *myb28/29* (Figure 4C). Next, we used a published dataset (Aubry et al., 2014) to test whether any of these gene sets were preferentially associated with the Arabidopsis bundle sheath. Genes down-regulated only in *myc2/3/4* (Figure 4D) or *myb28/*29 (Figure 4E) were not preferentially expressed in either the bundle sheath or whole leaf. However, of the 54 genes down-regulated in both *myc2/3/4* and *myb28/29* and present in the Aubry et al., (2014) dataset, 47 were strongly bundle sheath preferential (log2(bundle sheath/whole leaf) > 0.7) (Figure 4F). Only four genes were strongly depleted in the bundle sheath (log2(bundle sheath/whole leaf) < -0.7) (Figure 4F). Consistent with these down-regulated genes being directly regulated by the MYC-MYB system, motif enrichment analysis showed that the MYC2/MYC3/MYC4 (cluster 8) and MYB28/MYB29 (cluster 18) motifs were strongly enriched in promoters of genes down-regulated in both *myc2/3/4* and *myb28/29* mutants (Figure 4I). This was not the case for the *myc2/3/4* mutant (Figure 4G) or the *myb28/29* mutant alone (Figure 4H).

**Figure 4:**
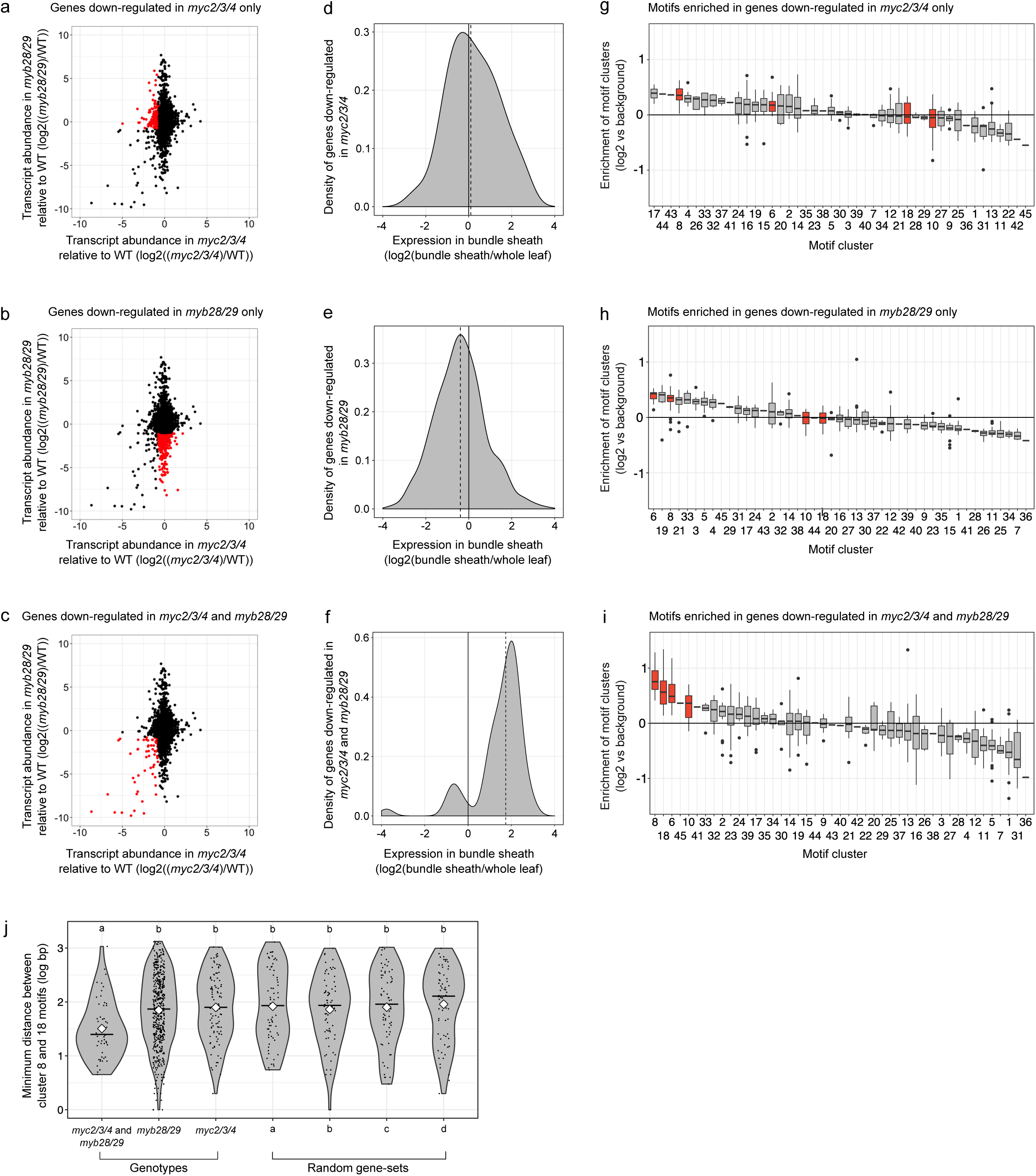
The MYC-MYB module controls bundle sheath expression of multiple genes. A, B, C) Change in expression in *myc2/3/4* compared with wild type (Major et al., 2017) plotted against that of *myb28/29* compared with wild type (Burow et al., 2015). Down-regulated genes (log2 < -0.75) in *myc2/3/4* only (A), *myb28/29* only (B) and in both *myc2/3/4* and *myb28/29* (C) are marked in red. D, E, F) Density plots for down-regulated genes highlighted in A, B and C indicating their expression in bundle sheath cells compared with whole leaves (Aubry et al., 2014). G, H, I) Enrichment analysis of motif clusters found in promoters of down-regulated genes highlighted in A, B and C. Clusters containing possible MYC binding sites (G-boxes) (Clusters 6 and 8) and MYB binding sites (Clusters 10 and 18) are highlighted in red. Note that Clusters 6, 8, 10 and 18 are strongly enriched in genes down-regulated in both *myc2/3/4* and *myb28/29*. Box-plots show inter-quartile range as upper and lower confines of the box, median as a solid black line and whiskers as maximum and minimum values excluding outliers. J) Violin plots depicting minimum distance (log bp) between cluster 8 and 18 motifs in promoters of genes highlighted in A, B, C and in four random sets of genes from Arabidopsis, ordered by median from smallest to largest. The median is shown as a horizontal black line, the mean as a white diamond. a and b indicate statistically significant differences (p < 0.05) determined by pairwise Wilcoxon rank-sum tests. n = individual genes tested. n=53 for genes down-regulated in *myc2/3/4* and *myb28/29*, n=506 for genes down=regulate only in *myb28/29*, n=101 in genes down-regulated only in *myc2/3/4* and n=66 for each of the random sets of Arabidopsis genes.

Previous work suggested that MYBs and MYCs bind to adjacent regions of promoters to activate expression of *GLS* genes (Schweizer et al., 2013). This was also the case in the *MYB76* DHS (Figure 3A). To test if this was also true for genes down-regulated in both *myc2/3/4* and *myb28/29* we investigated minimum distance between cluster 8 and 18 motifs in each set of promoters and compared this with random sets of genes. This showed that as well as being more enriched in the promoters of down-regulated genes in both mutants (Figure 4I) where cluster 8 and 18 motifs were both present, they were closer together in the down-regulated genes common to *myc2/3/4* and *myb28/29* compared with those down-regulated in only one mutant background, or in any of the random sets (Figure 4J). In summary, genes down-regulated in both *myc2/3/4* and *myb28/29* were generally strongly preferential to the bundle sheath, showed an enrichment of MYC2/3/4 and MYB28/29/76 binding sites in their promoters and where these motifs are both present, they were closer together than the other gene sets that we assessed. Taken together these findings suggest that the MYC-MYB module is important in controlling the expression of at least forty-seven genes in the bundle sheath of Arabidopsis.

## Discussion

### A positive regulator patterning gene expression to the bundle sheath

Although many promoters allowing expression in defined tissues of the shoot or root have been reported, our understanding of how these expression domains are generated is limited. Examples include promoters that drive expression in tissues such as apical meristems (Baima et al., 1995; Levin and Meyerowitz, 1995; Fletcher et al., 1999; Otsuga et al., 2001; Sarkar et al., 2007) the stele (Helariutta et al., 2000; Bonke et al., 2003), endodermis (Wysocka-Diller et al., 2000), cortex (Heidstra et al., 2004; Mustroph et al., 2009) and trichoblasts or atrichoblasts (Ruzicka et al., 2007; Masucci et al., 1996) of the root, as well as guard cells (Nakamura et al., 1995), phloem companion cells (Imlau et al., 1999) and epidermal cells (Thoma et al., 1994) of the shoot. In leaves, perhaps the best characterised of these promoters come from analysis of C_4_ species (Gowik et al., 2004; Akyildiz et al., 2007; Wiludda et al., 2012). For example, mesophyll specific expression of *PPCA1* from *F. trinervia* is due to two modules in a distal region of the promoter (Gowik et al., 2004; Akyildiz et al., 2007), whilst the *GLDPA* promoter from C_4_ *F. bidentis* generates expression in the bundle sheath because proximal promoter sequence leads to transcripts derived from a distal promoter element being degraded in mesophyll cells (Engelmann et al., 2008; Wiludda et al., 2012). In roots, using *SHORTROOT* as a model, it has been proposed that multiple *cis*-elements recognised by a complex network of both activators and repressors confine expression to the root vasculature (Sparks et al., 2016). Analysis of transcription factor binding *in vivo* is consistent with a highly combinatorial mosaic of regulatory DNA underpinning patterns of gene expression (Neph et al., 2012). To our knowledge, and perhaps due to this highly complex regulatory landscape, there are no examples of *cis*-elements and interacting transcription factors that limit gene expression to specific cell types in leaves. In this work, we show that combining DNaseI-SEQ data with functional analysis allowed identification of *cis-*elements and cognate transcription factors that pattern expression to bundle sheath cells of the leaf.

The module that generates expression in bundle sheath cells contains two *cis*-elements recognised by MYC and MYB transcription factors. When sequence containing this region is oligomerised it leads to strong and specific expression in the bundle sheath and so represents a short sequence that could be used to mis-express genes in this tissue. Bundle sheath cells link the vasculature to the photosynthetic mesophyll cells. In C_3_ species, they play important roles including the control of hydraulic conductance (Sage, 2001; Shatil-Cohen et al., 2011, Griffiths et al., 2013), transport of metabolites in and out of veins (Leegood, 2008), responses to high light episodes (Fryer et al., 2003), and assimilation of sulphur (Aubry et al., 2014). However, there are relatively few promoters available to drive or perturb expression in these cells (Takahashi et al., 2000; Wysocka-Diller et al., 2000). Short synthetic promoters have a number of advantages over the long promoter fragments currently available to direct gene expression to bundle sheath cells. These include reducing the likelihood of homology-based gene silencing if used more than once in any construct (Bhullar et al., 2003) and decreasing the chances of leakiness or off-target gene expression associated with use of full-length promoter fragments (Hernandez-Garcia & Finer, 2014). Oligomerization of *cis*-elements to achieve higher expression levels is a common strategy when creating synthetic promoters (Dey et al., 2015). As the *MYB76* DHS is short and can be oligomerized to tune expression levels, it appears to represent a promising fragment with which to perturb and modify functions including the control of hydraulic conductance, metabolite transport, responses to high light, and assimilation of sulphur in the bundle sheath.

### Control of bundle sheath expression by a MYC-MYB module

Glucosinolates are a diverse group of nitrogen and sulphur containing secondary metabolites that accumulate preferentially around the mid-vein and outer lamina in Arabidopsis rosette leaves and which are involved in defence against herbivory (Shroff et al., 2008). The methionine-derived aliphatic glucosinolate biosynthetic pathway is largely controlled by MYB28, MYB29 and MYB76 in combination with MYC2, MYC3 and MYC4 (Sønderby et al., 2007; Gigolashvili et al., 2008; Malitsky et al., 2008; Sønderby et al., 2010; Schweizer et al., 2013; Li et al., 2014). The *MYB76* promoter has previously been reported to generate expression in the vasculature (Gigoyashvili et al., 2008). Our analysis now shows that its expression domain includes the bundle sheath, and that this is under control of a MYC-MYB module. None of the MYC2, MYC3, MYC4, MYB28, MYB29 or MYB76 transcription factors were identified in our Yeast One-Hybrid analysis (Figure 3B) or previously published analysis of the whole *MYB76* promoter (Li et al., 2014). This may be due to these transcription factors requiring additional partners to bind DNA, consistent with the model of MYC-MYB dimers being required for the activation of target genes (Schweizer et al., 2013). Although this appears inconsistent with the *trans*-activation assays where infiltration of individual MYC transcription factors activated expression from the *MYB76* DHS (Figure 3C), this may be due to their interaction with endogenous MYBs from *N. benthamiana*. Alternatively, and consistent with the MYC binding site alone being able to generate weak expression in the bundle sheath (Figure 2D), the interaction may be below the detection limit of the Yeast One-Hybrid assay.

It is theoretically possible that this MYC-MYB module acts as a general activator of transcription across all cells of the leaf and another distinct mechanism represses expression in other cell types. However, for the following reasons, we favour a model in which the MYC-MYB module activates expression preferentially in the bundle sheath. First, the 256bp DHS which contains the MYC-MYB binding sites is sufficient for bundle sheath preferential expression, and the 27bp region containing the potential MYC binding site, but lacking MYB binding sites, is sufficient for weak expression in the bundle sheath. Second, *MYB28* and *MYB29* are preferentially expressed in the bundle sheath (Figure 1B) consistent with them activating expression in this cell-type. Third, other transcription factor binding sites required for directing bundle sheath preferential expression would have to be present in the 27bp region. This is possible, however we would expect such binding sites to be enriched in promoters of additional genes that are strongly expressed in the bundle sheath of wild-type plants but down regulated in both *myc2/3/4* and *myb28/29* mutants. No such binding sites were detected (Figure 4I). In summary, we cannot completely rule out a requirement for other factors operating in other cell types such that *MYB76* expression is restricted to the bundle sheath. However, the MYC-MYB module activating expression preferentially in the bundle sheath is a more parsimonious mechanism. To our knowledge, this MYC-MYB module provides the first example of a regulatory system governing the spatial control of gene expression in leaves.

At this point, how the MYC-MYB module directs preferential expression in the bundle sheath is not clear. *MYB28* and *29* transcripts accumulate preferentially in the bundle sheath (Figure 1B) but this is not as apparent for transcripts of genes encoding *MYC2, MYC3* and *MYC4* (Figure 3B). MYC transcription factors are regulated by jasmonic acid (JA) signalling thorough interactions with JASMONATE-ZIM DOMAIN (JAZ) proteins (Fernandez-Calvo et al., 2011; Howe et al., 2018). Whether there is a link between JA signalling and bundle sheath preferential gene expression remains to be determined.

### Roles of additional transcription factors in controlling *MYB76*

Many additional transcription factors were identified as binding the *MYB76* DHS in Yeast One-Hybrid analysis (Figure 3B) and some of these have been reported to bind the entire promoter previously (Li et al., 2014). These transcription factors are likely important for controlling other aspects of *MYB76* expression in addition to the spatial patterning determined by the MYC-MYB module. For example, multiple ERF family transcription factors interacted with the *MYB76* DHS (Figure 3B) and there is an ERF family transcription factor binding site in the DHS (Figure 3A). Additionally, DREB2A weakly activates expression from the *MYB76* DHS (Figure 3C) and *MYB76* transcripts were less abundant in a *dreb2a* mutant allele (Figure 3D). Recent work has linked auxin signalling and glucosinolate biosynthesis under drought conditions with DREB2A/B signalling (Salehin et al., 2019). This suggests that DREB2A may have a role in regulating *MYB76* expression in response to environmental stimuli. Because DREB2A transcripts do not accumulate preferentially in the bundle sheath (Figure 3B) it is not clear how activation of *MYB76* outside of the bundle sheath is avoided. Possibilities include DREB2A activity being limited to the bundle sheath by post-transcriptional and/or post-translational mechanisms or a requirement for other binding partners. Post-transcriptional and post-translational regulation has been reported for DREB2A (Agarwal et al., 2017) with post-transcriptional regulation by alternative splicing being reported (Egawa et al., 2006; Qin et al., 2007; Matsukura et al., 2010; Vainonen et al. 2012). Whilst overexpression of *DREB2A* in Arabidopsis does not affect the expression of target genes (Liu et al., 1998) an isoform lacking key phosphorylation sites activates the majority of DREB2A targets (Sakuma et al., 2006). Thus, post-translational regulation is essential for DREB2A activity suggesting a mechanism for restricting DREB2A activation of *MYB76* expression to the bundle sheath. The *MYB76* DHS also contains a MADS domain transcription factor binding site (Figure 3A). MADS domain transcription factors are involved in controlling gene expression required for flower development (Theißen et al., 2016) and could potentially be involved in directing *MYB76* expression to flowers where significant levels of GLS accumulate (Sarsby et al., 2012). In summary, as well as the MYC-MYB module generating bundle sheath preferential expression in leaves, multiple other transcription factor binding sites in the DHS may be important for controlling other aspects of *MYB76* expression such as responses to environmental stimuli and expression in different organs.

### The role of the MYC-MYB module outside glucosinolate biosynthesis

Our data are consistent with the MYC-MYB module patterning the expression of at least forty-seven genes to the Arabidopsis bundle sheath. This represents about 3.6% of the 1316 genes previously reported to be preferentially expressed in the bundle sheath (using a log2(bundle sheath/whole leaf) cut off >1) (Aubry et al., 2014) indicating that this module must operate alongside other networks. This notion is supported by previous analysis of promoters controlling bundle sheath preferential expression in Arabidopsis. For example the region identified as controlling the bundle sheath preferential expression of *SULTR2;2* (Kirschner et al., 2018) does not contain the MYC-MYB module.

Although extensive glucosinolate biosynthesis is confined to the Brassicaceae (Halkier & Gershenzon, 2006) there are indications that this MYC-MYB module patterns genes unrelated to glucosinolates to the bundle sheath. One possibility is associated with glucosinolate biosynthesis representing a derived pathway that has evolved in the Brassicaceae. It would seem more parsimonious if its patterning to bundle sheath cells was mediated by integration into an existing gene regulatory network associated with this cell type, than through evolution of a network *de novo*. This seems plausible because in addition to several enzymes of primary sulphur metabolism being part of the core glucosinolate biosynthesis pathway (Yatusevich et al., 2010), transcripts encoding many enzymes of sulphur transport and assimilation are more abundant in the bundle sheath compared with whole leaves (Aubry et al., 2014). This raises the possibility that compartmentation of glucosinolate biosynthesis to the bundle sheath may have occurred through acquisition of *cis*-elements that restrict the expression of sulphur assimilation genes to this cell-type.

In addition to genes associated with sulphur metabolism, there is also evidence that this MYC-MYB module may pattern genes that are thought to represent some of the first steps towards evolving C_4_ photosynthesis. Although the Arabidopsis *GLYCINE DECARBOXYLASE P-PROTEIN 1 (GLDP1*) gene is expressed strongly in both the mesophyll cells and the vasculature, deletion of an M-box in the promoter resulted in bundle sheath expression of *GLDP1* (Adwy et al., 2015). The remnant expression in the bundle sheath and vein tissue was associated with a 266bp region named the V-box (Adwy et al., 2015). Re-analysis of these 266bp identified MYC and MYB binding sites within 25bp of each other (Supplemental Figure 12). It is possible that constitutive expression of *GLDP1* in the C_3_ Arabidopsis leaf is due to the M-box and MYC-MYB modules driving expression in mesophyll and bundle sheath strands respectively. This is consistent with the MYC-MYB module being an activator of expression in the bundle sheath and therefore its presence not preventing activation in other cell types. This would explain why only some genes containing the MYC-MYB module are preferentially expressed in bundle sheath strands and suggests that bundle sheath preferential expression is partly defined by lack of activation in other cell-types.

One of the early events associated with the transition from C_3_ to C_4_ photosynthesis is thought to be the restriction of the Glycine Decarboxylase complex to the bundle sheath as part of establishing a C_2_ photosynthetic cycle (Mallmann et al., 2014). The M-box of *GLDP1* is highly conserved in the Brassicaceae, but is lost in *Moricandia nitens*, a species that uses C_2_ photosynthesis and partitions *GLDP* to bundle sheath cells. Conversely the V-box, and predicted MYC and MYB binding sites, is conserved in *M. nitens* (Adwy et al., 2015; Adwy et al., 2019). It is therefore possible that during the evolution of C_2_ photosynthesis in the Brassicaceae, the MYC-MYB module in *GLDP* is responsible for bundle sheath expression of the *GLDP* gene once the M-box is lost.

In summary we report a MYC-MYB module that directs gene expression to the bundle sheath of Arabidopsis. To our knowledge, this provides the first example of a regulatory system governing the spatial control of gene expression in leaves. In the future it will be interesting to determine if this module has been co-opted during the evolution of C_4_ photosynthesis to pattern components of the C_4_ cycle to this cell type.

## Materials and methods

### Plant material, growth conditions and cloning

Seed of Arabidopsis was sterilised by washing in 70% (v/v) ethanol for 3 minutes followed by washing in 100% ethanol for 1 minute. Transformants were selected on 0.5% (w/v) Murashige & Skoog medium (pH 5.8) 1% (w/v) agar with the relevant antibiotics. After 2-3 days of stratification in the dark at 4°C, tissue culture plates were transferred to a 16 hour photoperiod growth chamber with a light intensity of 200 μmol m^-2^ s^-1^ photon flux density, 65% relative humidity and a temperature cycle of 24°C (day) and 20°C (night). Transformed seedlings were transferred onto 3:1 Levington M3 high nutrient compost and Sinclair fine Vermiculite soil mixture and grown for another 2-3 weeks before analysis. *N. benthamiana* plants used for transient assays were grown from seed in pots containing the same soil mixture with a 16 hour photoperiod, 200 µmol m^-2^ s^-1^ photon flux density, 60% relative humidity and 22°C.

T-DNA insertion mutants for *dreb2a* (GK-179C04) were obtained from the Nottingham Arabidopsis Stock Centre (NASC). T-DNA insertion lines were genotyped to identify lines homozygous for the required T-DNA insertion and RT-PCR was performed to confirm that the mutation resulted in the loss of *DREB2A* gene expression.

The full length *MYB76* gene as well as the promoter alone were amplified from Arabidopsis *Col-0* genomic DNA and then fused to *uidA*. The minimal CaMV35S promoter was synthesised and fused to *MYB76 DH*S by polymerase chain reactions (PCR). Deletion of the DHS within the promoter was achieved by PCR fusion of the 5’ end of the promoter with the 3’ end of the promoter prior to being cloned into the pENTR/D TOPO vector. Each forward primer contained a CACC overhang to ensure directional cloning. A Gateway LR reaction was performed to transfer the relevant inserts into a modified pGWB3 vector (Nakagawa et al., 2007) that contained an intron within the *uidA* sequence. *MYB76gDNA::uidA*, *2xDHS_CaMV35SMin::uidA* and *2x27_CaMV35SMin::uidA* were constructed using Golden Gate technology (Weber et al., 2011). Motif substitutions were made using the QuikChange Lightning Site-Directed Mutagenesis (Agilent Technologies) and motif deletions were made by overlapping PCR. Constructs were then placed into *Agrobacterium tumefaciens* strain GV3101 and introduced into Arabidopsis Col-0 by floral dipping (Clough & Bent, 1998).

Constructs for *trans-*activation assays were made using the Golden Gate system. Coding sequence of candidate transcription factors were cloned from Arabidopsis cDNA, domesticated to remove *Bpi*I and *Bsa*I restriction sites and cloned into level 0 vectors. Level 1 constructs were generated to constitutively express candidate transcription factors, to constitutively express a p19 silencing suppressor, to constitutively express a GUS reporter to act as an infiltration control and to fuse the *MYB76* DHS with a *LUCIFERASE* reporter to provide an output of activation from the DHS. These level 1 constructs were then assembled into level 2 modules and transformed into *A. tumefaciens* GV3101. Constructs for the constitutively active LjUBI promoter, the OCS1 terminator, LUC coding region, GUS coding region have been published previously (Feike et al., 2019) and parts were cloned into appropriate Golden Gate vectors (Patron et al., 2015).

### GUS staining, MUG assays, and GFP imaging

To take into account position effects associated with transgene insertion site, GUS staining was undertaken on at least six randomly selected T1 plants for each *uidA* fusion (Jefferson et al., 1987). The staining solution contained 0.1 M Na_2_HPO_4_ pH 7.0, 2 mM Potassium ferricyanide, 2 mM Potassium ferrocyanide, 10 mM EDTA pH 8.0, 0.06% (v/v) Triton X-100 and 0.5 mg ml^-1^ X-gluc. Leaves from three-week old plants were vacuum-infiltrated three times in GUS solution for one minute and then incubated at 37°C for between 3 and 72 hrs depending on the strength of the promoter being assessed. Next, stained samples were fixed in 3:1 (v/v) ethanol:acetic acid for 30 minutes at room temperature, cleared in 70% (v/v) ethanol at 37°C and then placed in 5 M NaOH for 2 hrs. Samples were stored in 70% (v/v) ethanol at 4°C. To quantify reporter accumulation from each promoter the quantitative assay that assesses the rate of MUG conversion to 4-methylumbelliferone (MU) was performed (Jefferson et al., 1987) on between 10 and 25 lines. Tissue was frozen in liquid nitrogen, homogenised and soluble protein extracted in 5 volumes of Protein extraction buffer (1 mM MgCl_2_, 100 mM NaCl, 50 mM Tris (Melford) pH7.8). 15 μl of protein extract was incubated with 60 μl of MUG at 37 °C for one, two, three and four hours respectively. The reaction was stopped after each time point by addition of 75 μl 200 mM anhydrous sodium carbonate. GUS activity was analysed via measurements of fluorescence of MU at 455 nm after excitation at 365 nm. The concentration of MU/unit fluorescence in each sample was interpolated using a concentration gradient of MU over a linear range.

GFP imaging was performed on at least seven independent T1 lines of *pMYB76::H2B::GFP* and *2xMYB76_DHS::H2B::GFP.* Rosette leaves of four week old plants were sampled and outer tissue layers were removed by scraping leaves with a razor blade under 1X PBS solution. Samples were imaged on a Leica TCS SP8 confocal microscope, GFP was excited at 488 nm and emission was detected at 500-530 nm. Images were recorded on LAS Image analysis software (Leica) and processed to merge channels and add scale bars in ImageJ v1.52a (https://imagej.nih.gov/ij/).

### Yeast One-Hybrid screen

Regions screened for transcription factor binding via Yeast One-Hybrid were first inserted into pENTR 5’TOPO TA entry vector (Thermofisher) and subsequently placed into the pMW2 and pMW3 destination vectors containing *HIS3* and *LACZ* marker genes respectively (Deplancke et al., 2006). The enhanced Yeast One-Hybrid screen against a complete collection of 2000 Arabidopsis transcription factors was undertaken as described previously (Gaudinier et al., 2011; Pruneda-Paz et al., 2014; Gaudinier et al., 2017). Details of the bait sequence and list of interactors found in Supplemental table 4.

### *Trans* activation assays and qRT-PCR

To test interactions between the *MYB76* DHS and transcription factors *in planta* transient infiltration of *N. benthamiana* was performed. Overnight cultures of *A. tumefaciens* were pelleted and re-suspended in infiltration buffer (10mM MES (pH5.6), 10mM MgCl_2_, 150μM acetosyringone) to an optical density of 0.3. Cultures were then incubated for 2hrs at room temperature and infiltrated into the abaxial side of leaves of four-week old plants with a 1 ml syringe.

Leaf discs from infiltrated regions were sampled 48hrs after infiltration and flash frozen in liquid N_2_. Protein for MUG and LUC assays was extracted on ice in 1x passive lysis buffer (PLB: Promega). MUG assays were performed by adding 40μl of protein extract to 100μl of MUG assay buffer (2mM MUG, 50mM Na_3_PO_4_/Na_2_PO_4_ buffer (pH 7.0), 10 mM EDTA, 0.1% (v/v) Triton X-1000, 0.1% (w/v) Sodium Lauroyl sarcosinate and 10 mM DTT). Stop buffer (200 mM Na_2_CO_3_) was added at 0 and 30 mins and rate of MUG accumulation was measured in triplicate on a plate reader (CLARIOstar, BMG lab tech) with excitation at 360 nm and emission at 465 nm. LUC activity was measured with 20 μl of protein sample and 100 μl of LUC assay reagent (Promega). Activation from the DHS was calculated as (LUC luminescence/rate of MUG accumulation) * 100.

Single rosette leaves from four-week old col-0, *dreb2a* and *myc2/3/4* plants were sampled six hours after the onset of light and flash frozen in liquid N_2_. RNA was extracted with the RNeasy plant mini-kit (Qiagen) as manufacturer’s instructions and cDNA was synthesised using the Superscript double stranded cDNA synthesis kit as manufacturer’s instructions (Invitrogen) with on-column DNase1 treatment. qRT-PCR was performed using SYBR green master mix (Bio-Rad) on a CFX384 touch Real-Time PCR Detection System (Bio-RAD). Transcripts of *MYB76* were normalised to the expression of *ASCORBATE PEROXIDASE 3 (APX3: At4g35000), ASPARTIC PROTEINASE A1 (APA1: At1g11910)* and *UBIQUITIN CONJUGATING ENZYME 21 (UBC21: AT5G25760).* Relative expression was determined using the single delta Ct method and the data reported are from normalisation against *APA1.* Results were very similar regardless of the reference genes used.

### Datasets, GO term analysis and *de novo* motif identification

Computational analysis used previously published Arabidopsis datasets for bundle sheath and whole leaf translatomes (Aubry et al., 2014), *myc2/3/4* mutants and col-0 (Major et al., 2017), and *myb28/29* mutants and col-0 (Burow et al., 2015). For Gene Ontology (GO) term enrichment analysis the top 200 most bundle sheath preferential (log2(bundle sheath expression/35S expression) genes in the cell-type specific translatome (Aubry et al., 2014) were used as input. The AgriGO tool (Du et al., 2010) was used with default parameters and all genes annotated in the Arabidopsis TAIR10 genome were sued as background. The MEME tool from The Multiple Em for Motif Elucidation (MEME) suite v.4.8.1. (Bailey et al., 2009) was applied to search for conserved motifs within promoter sequences of genes expressed in the Arabidopsis bundle sheath. Maximum length of the motif was set to eight nucleotides, both strands of the sequence were searched and each motif had to be present in every sequence.

To cluster transcription factor binding motifs the RSAT matrix-clustering tool (Castro-Mondrago et al., 2017) was run on all Arabidopsis motifs from the JASPAR motif database (Fornes et al., 2019) using default parameters which generated 43 motif clusters. The Find Individual Motif Occurances (FIMO) (Grant et al., 2011) was used to scan DNA sequences for matches to Arabidopsis transcription factor binding motifs found in the JASPAR motif database (Fornes et al., 2019). To account for input sequence composition a background model was generated using the fasta-get-markov tool from the MEME suite (Bailey et al., 2009). FIMO was then run with default parameters and a p-value cut-off of 1e-04.

Motif enrichment in promoters of gene sets was analysed using a custom BASH script. Promoter regions (1500bp) were extracted using the getfasta tool from Bedtools (Quinlan and Hall, 2010). These promoters were scanned for transcription factor binding motifs using FIMO (as above) and counts of motifs in gene sets were recorded. Frequency of a given motif in a gene set was calculated as a proportion of the total motifs and enrichment was calculated as frequency vs background frequency. Background frequency was defined as mean motif frequency in promoters of three random sets of 2000 Arabidopsis genes. Results of motif frequency analysis presented as the log2 of enrichment and motifs sorted by motif cluster. FIMO outputs were sorted to only include matches to cluster 8 and 18 motifs and a custom Python script was used to find the minimum distance between the centres of cluster 8 and 18 motif in the same promoter.

### Statistical analysis

For statistical analysis extreme outliers were identified and removed from analysis. Normality of the data was assessed using the Shapiro-Wilks test. Where data were normally distributed pairwise T-tests were used to assess significance. Where data were not normally distributed, Wilcoxon rank-sum tests were used to assess significance. Levene’s test was used to assess equality of variance. Where variance was equal standard deviations were pooled, where variance was not equal variance was not pooled. All tests were two-sided. Pairwise T-tests with pooled SD were used to assess significance in *trans-*activation assays and without pooled SD in qRT-PCR assays. Wilcoxon rank sum tests were used to assess significance for differences in distributions of minimum differences between cluster 8 and 18 motifs in different gene sets. All statistical analysis was performed using R (RStudio Team, 2015) and plots generated using ggplot2 (Wickham, 2009).

### Code availability

All code associated with this manuscript is available in the Github repository: https://github.com/hibberdlab/Knerova_Dickinson_Arabidopsis_bundle_sheath_bipartite_module.

### Data availability

Underlying data required to generate plots are available in the Github repository: https://github.com/hibberdlab/Knerova_Dickinson_Arabidopsis_bundle_sheath_bipartite_module. All other data available on request.

---

**Supplemental Figure 1:**
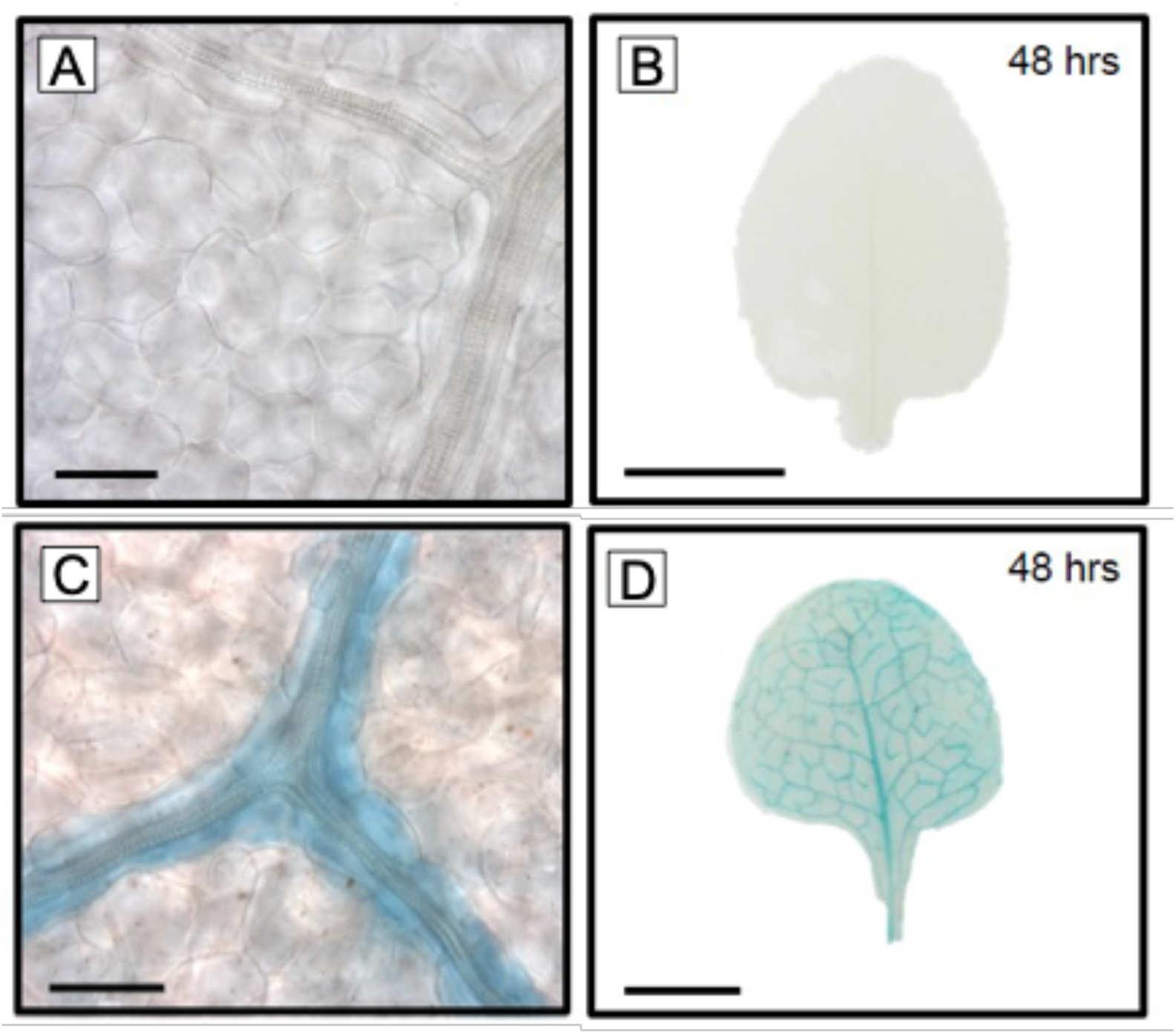
Representative images of *proMYB*28 and *proMYB29* GUS. Staining performed for 48hrs and scale bars represent 0.5 cm (a and c) and 50 µm (b and d).

**Supplemental Figure 2:**
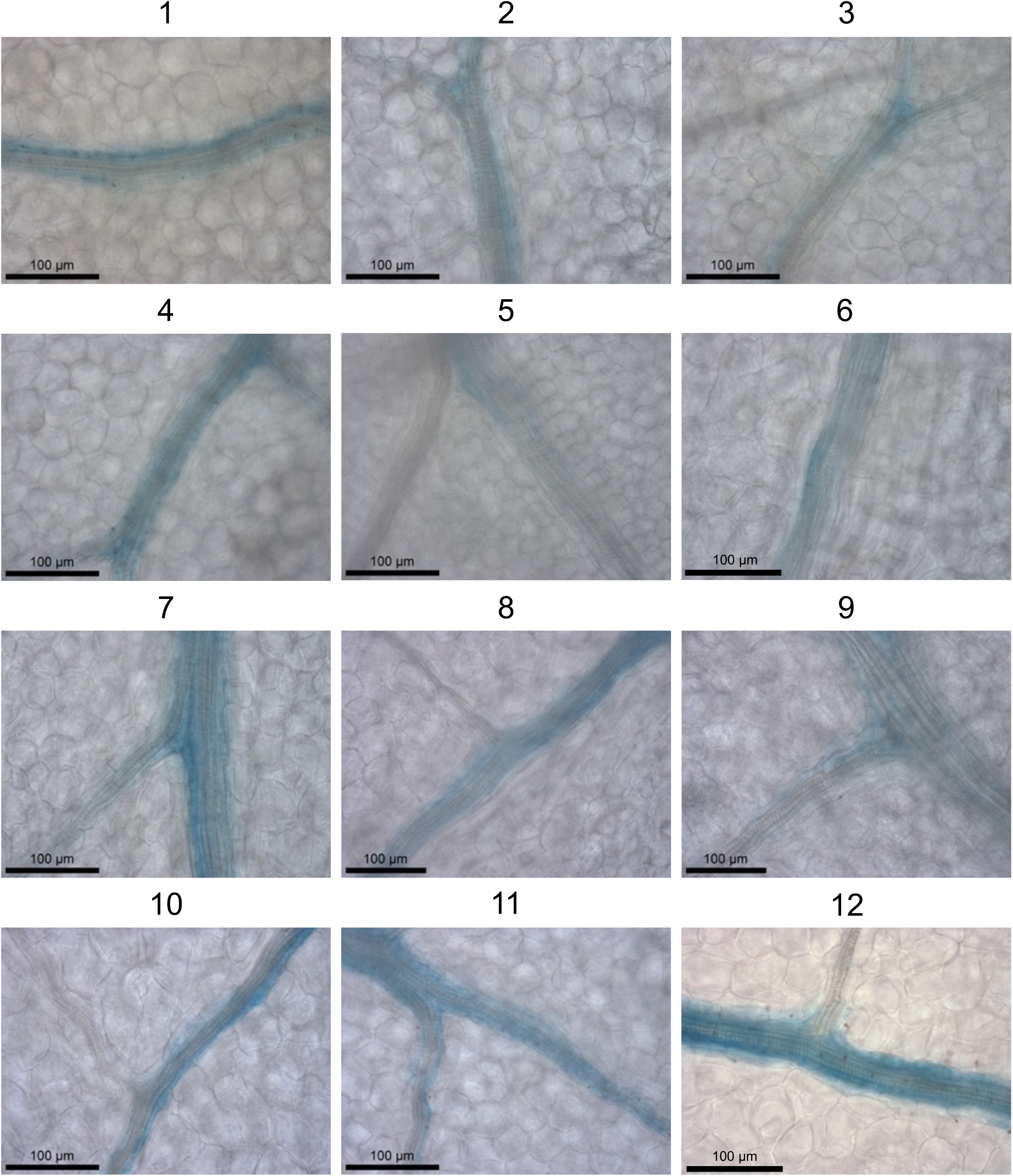
Nucleotides -1725 to +279 relative to the predicted translational start site of *MYB76* generate preferential expression in the bundle sheath. Images from twelve independent transgenic lines. Leaves were stained for 30hrs. Scale bars represent 100 µm.

**Supplemental Figure 3:**
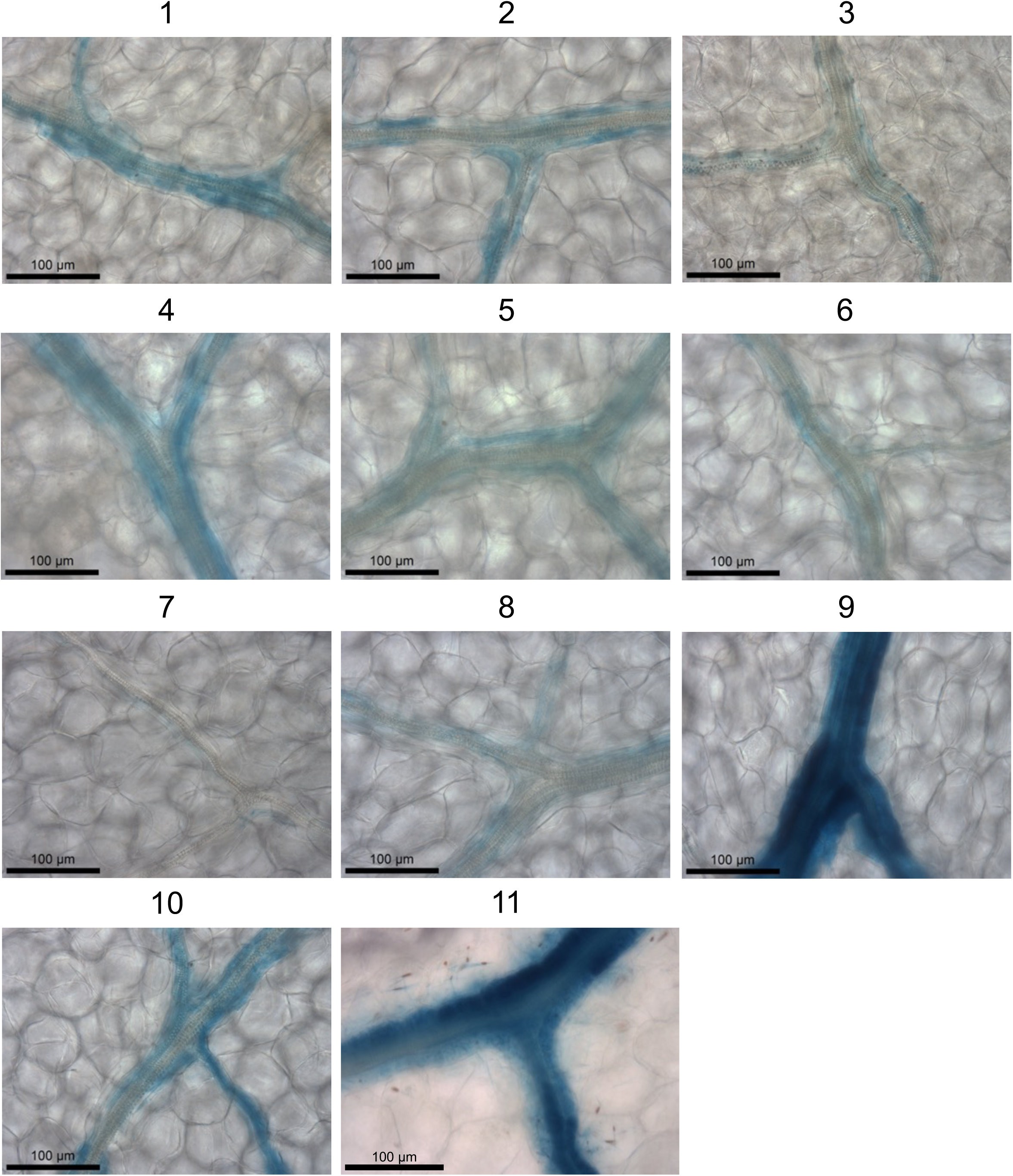
The genomic *MYB76* sequence fused to GUS generates preferential expression in the bundle sheath. Images from eleven independent transgenic lines Leaves were stained for 72hrs except line 11 which was stained for 48hrs. Scale bars represent 100 µm.

**Supplemental Figure 4:**
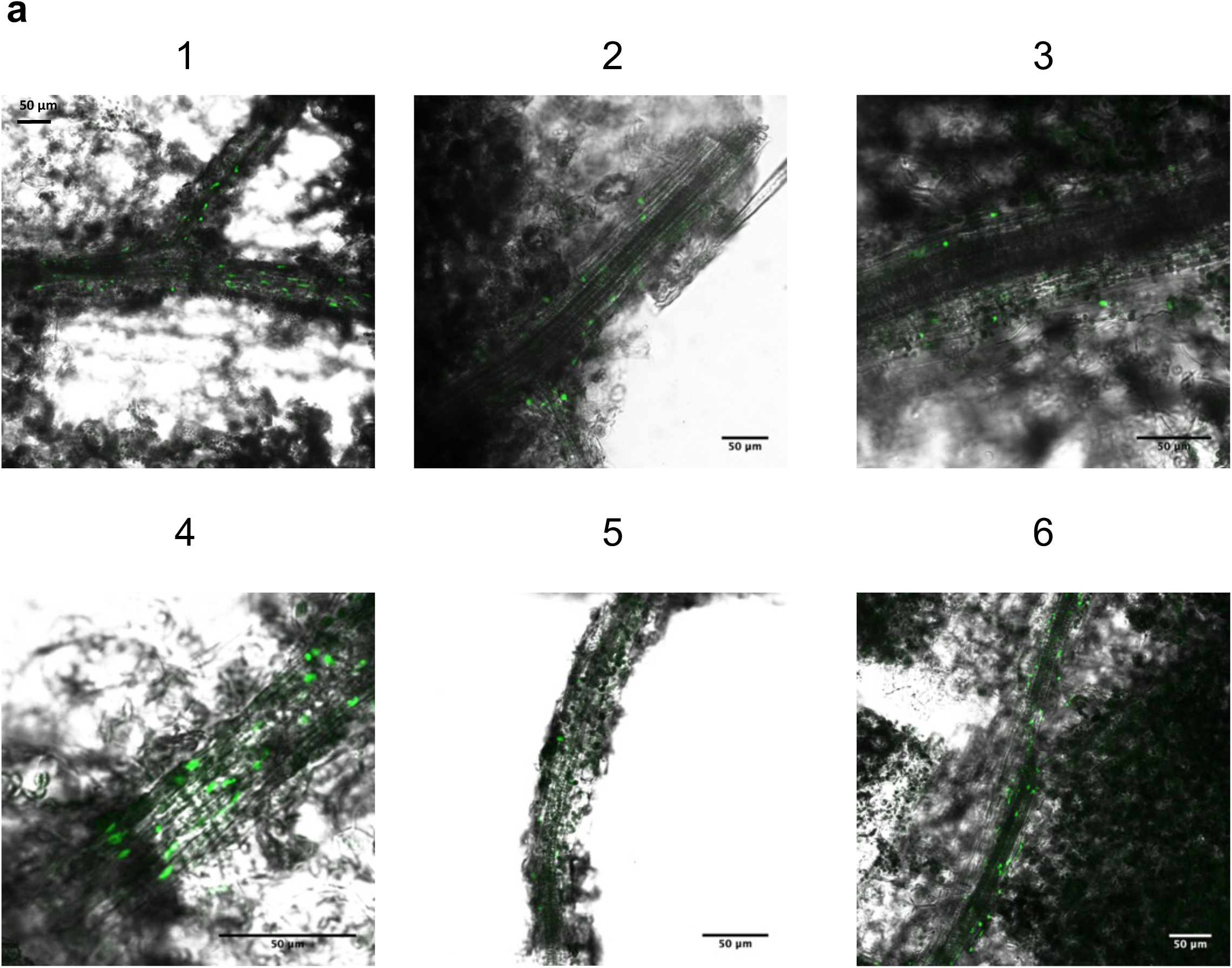
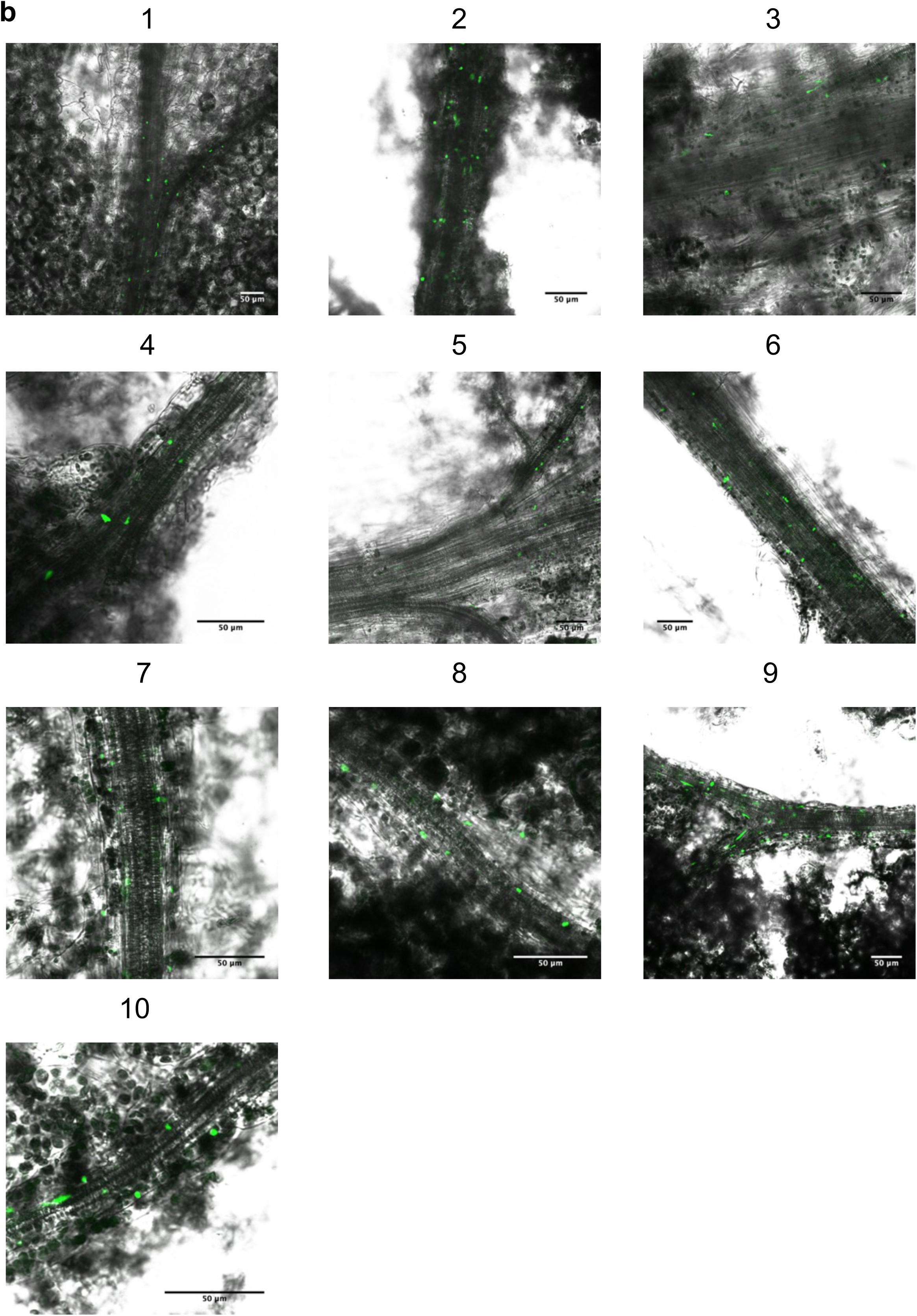
a) Representative images from independent T1 lines of the *MYB76* promoter driving expression of a histone GFP fusion *(H2B::GFP).* b) Representative images from independent T1 lines of 2x the *MYB76* DHS driving expression of a histone GFP fusion *(H2B::GFP)*.

**Supplemental Figure 5:**
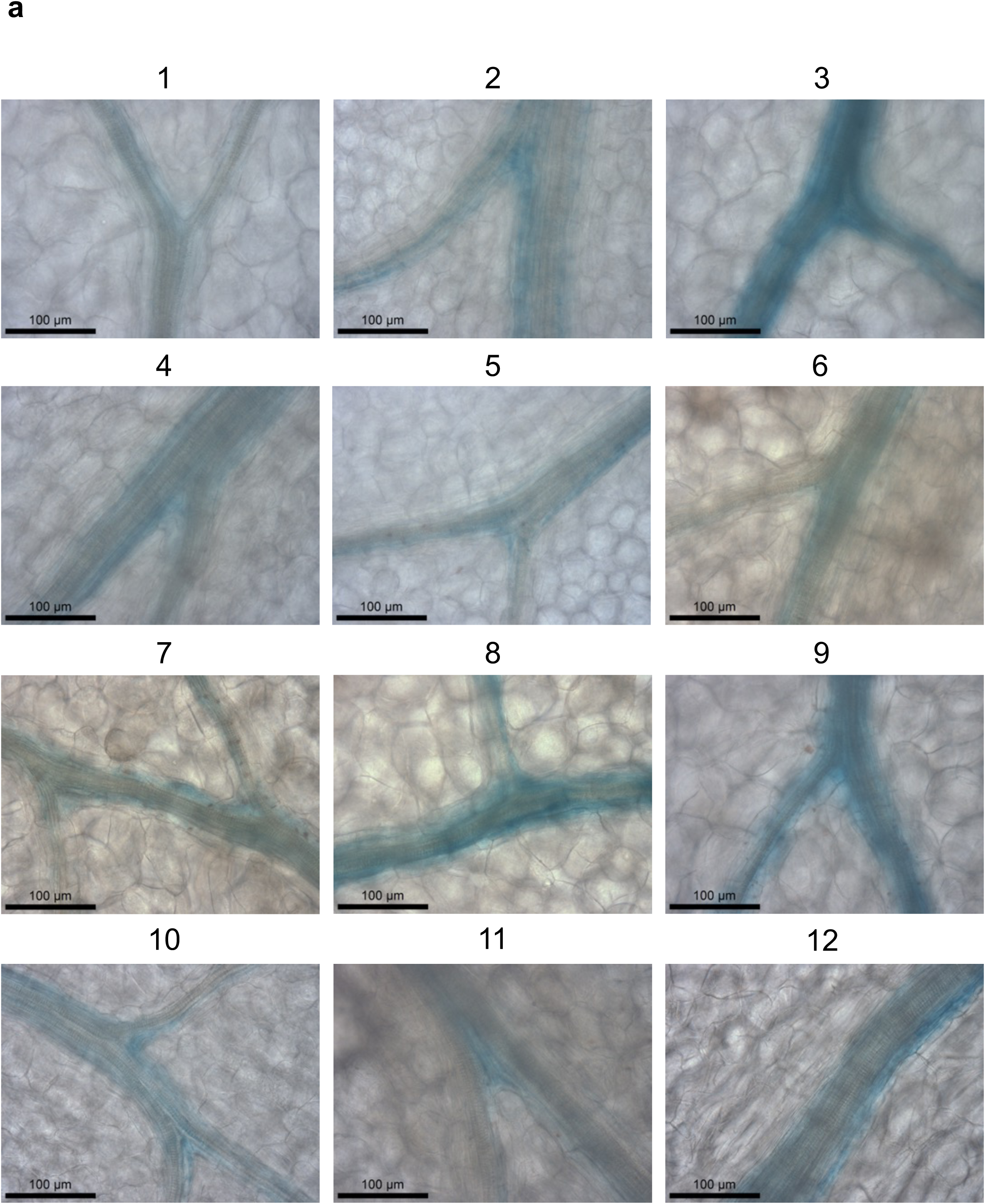
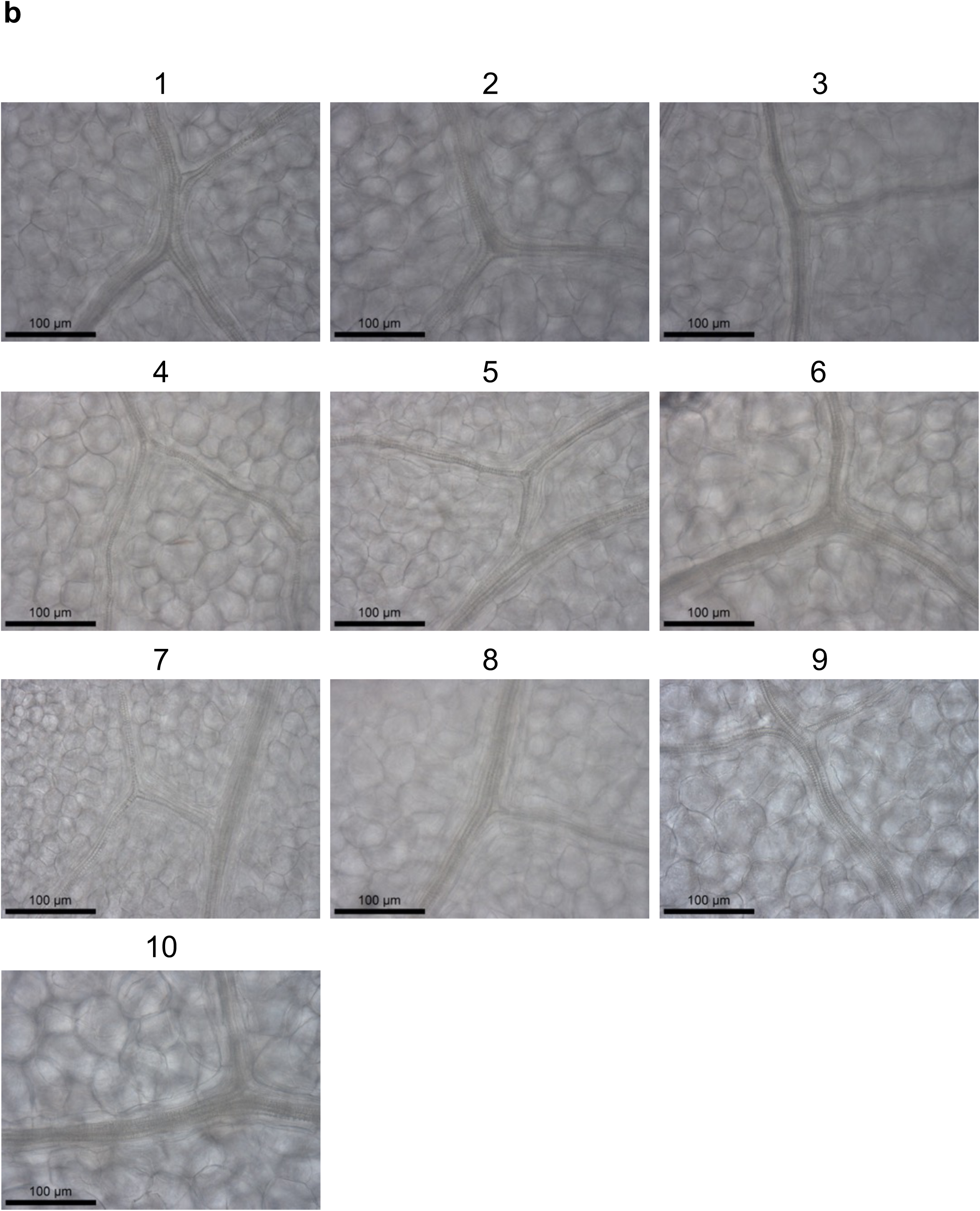
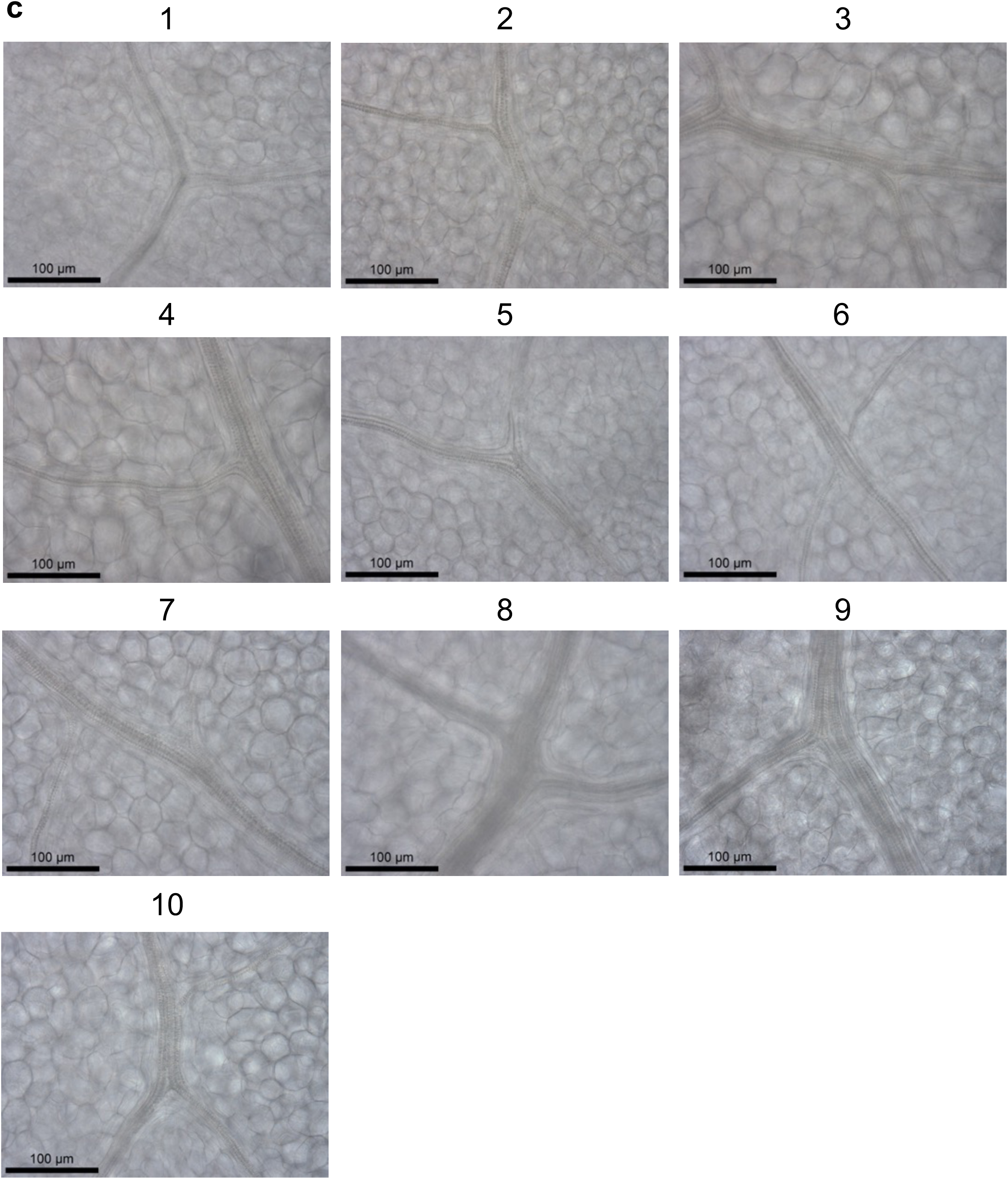
a) Nucleotides -1264 to +279 relative to the predicted translational start site of MYB76 generate preferential expression in the bundle sheath. Images from twelve independent transgenic lines. b) 796bp of the promoter combined with the first exon and intron of *MYB76* does not generate BS preferential expression. Images from ten independent transgenic lines. c) 294bp of the promoter combined with the first exon and intron of *MYB76* does not generate BS preferential expression. Images from ten independent transgenic lines. Leaves were stained for 48hrs. Scale bars represent 100 µm.

**Supplemental Figure 6:**
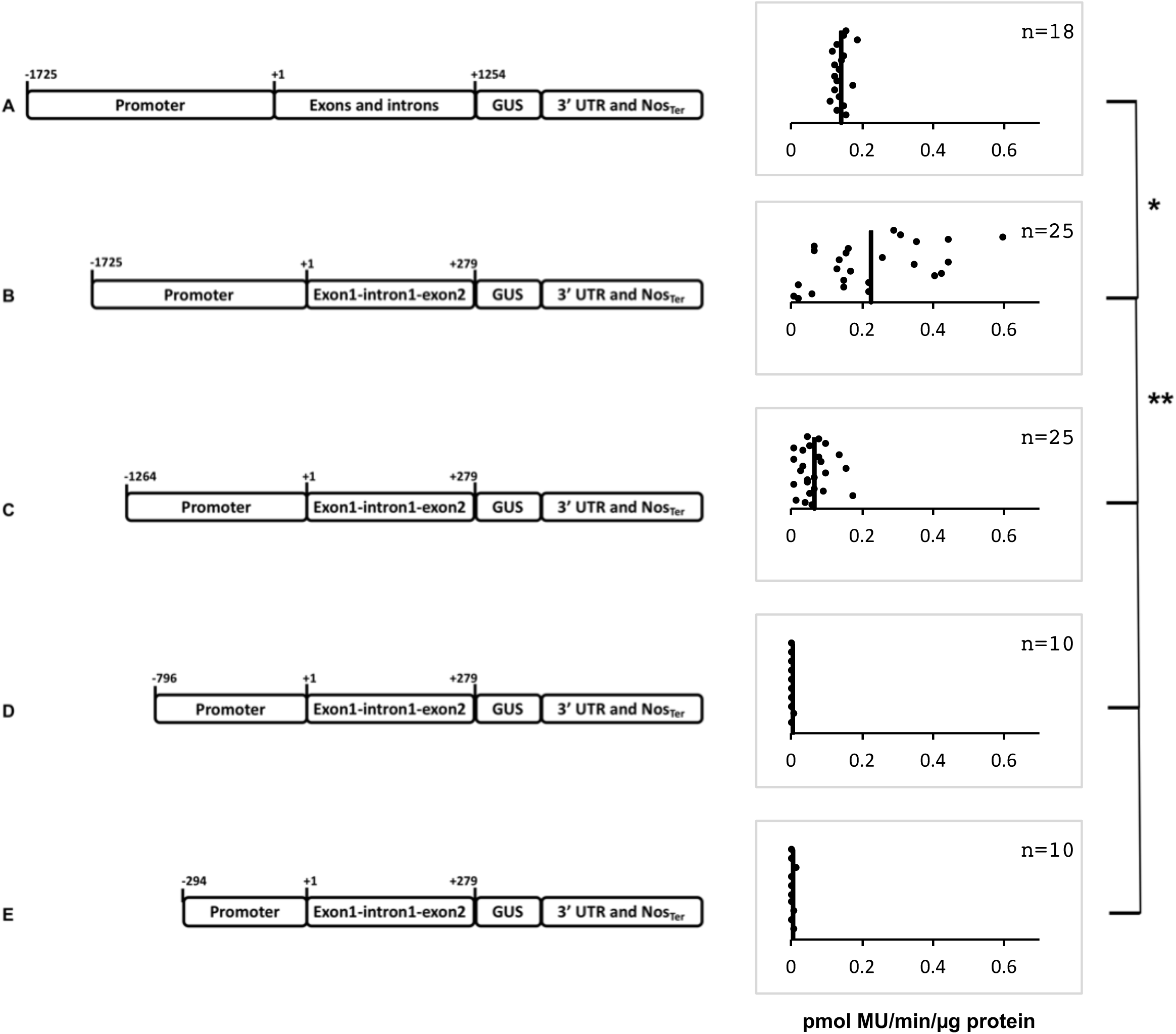
Schematics of deletion series (left) and quantification of MUG activity via the flourometric assay (right) for multiple independent transformants from each construct. (A) The *MYB76* genomic sequence. (B) The *MYB76* promoter including the first two exons and the first intron fused to the *uidA*. (C-E) A region between -1264 and -796bp is responsible for GUS accumulation in the bundle sheath. The fluorometric MUG assay shows quantitative repressors and enhancers are located in the gene and in the promoter respectively. X-axis indicates GUS activity and individual biological replicates are ordered randomly on the y axis. n, number of multiple independent transformants; *, p value < 0.05; ** p value < 0.001 determined by T-tests.

**Supplemental Figure 7:**
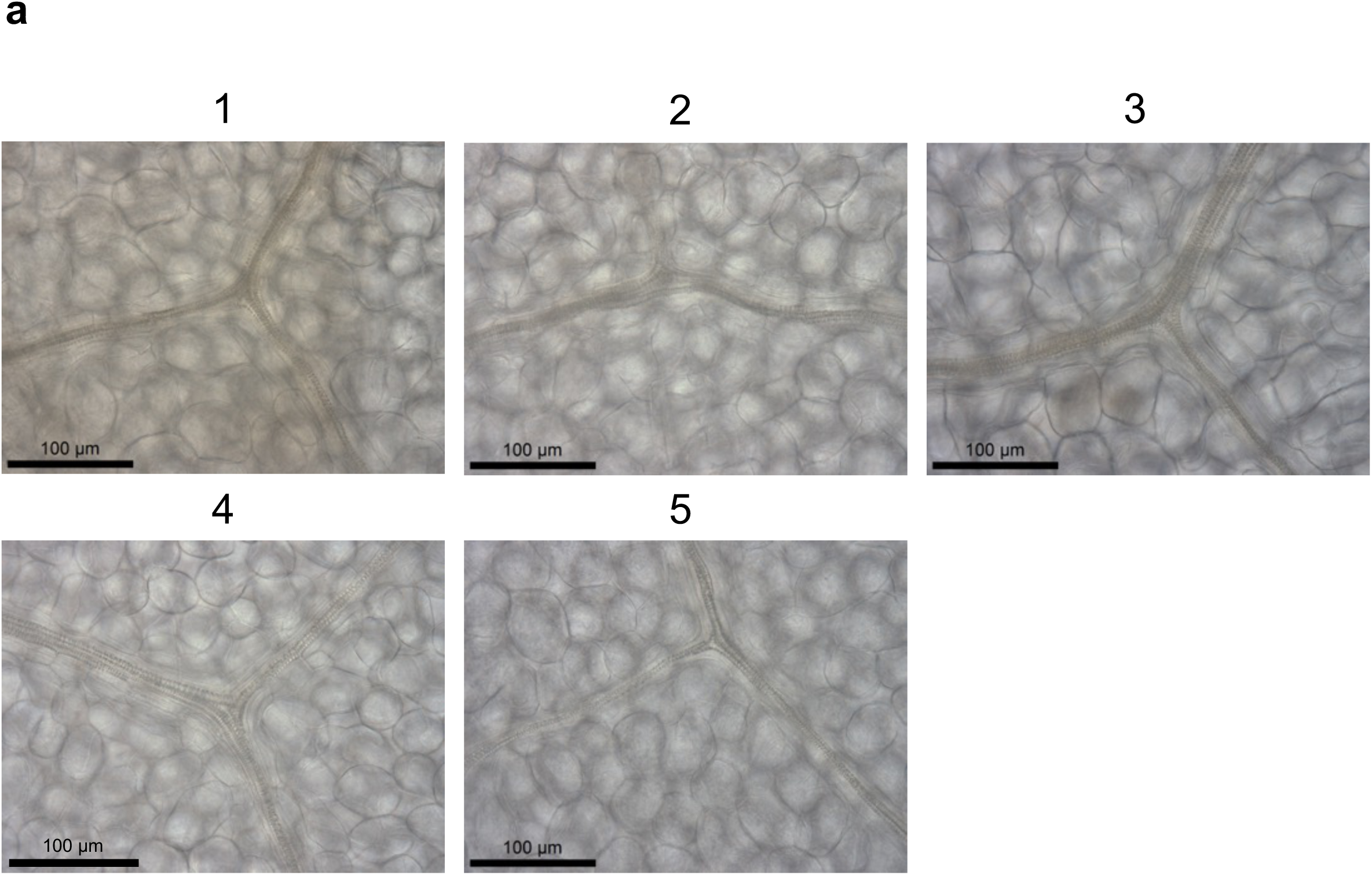
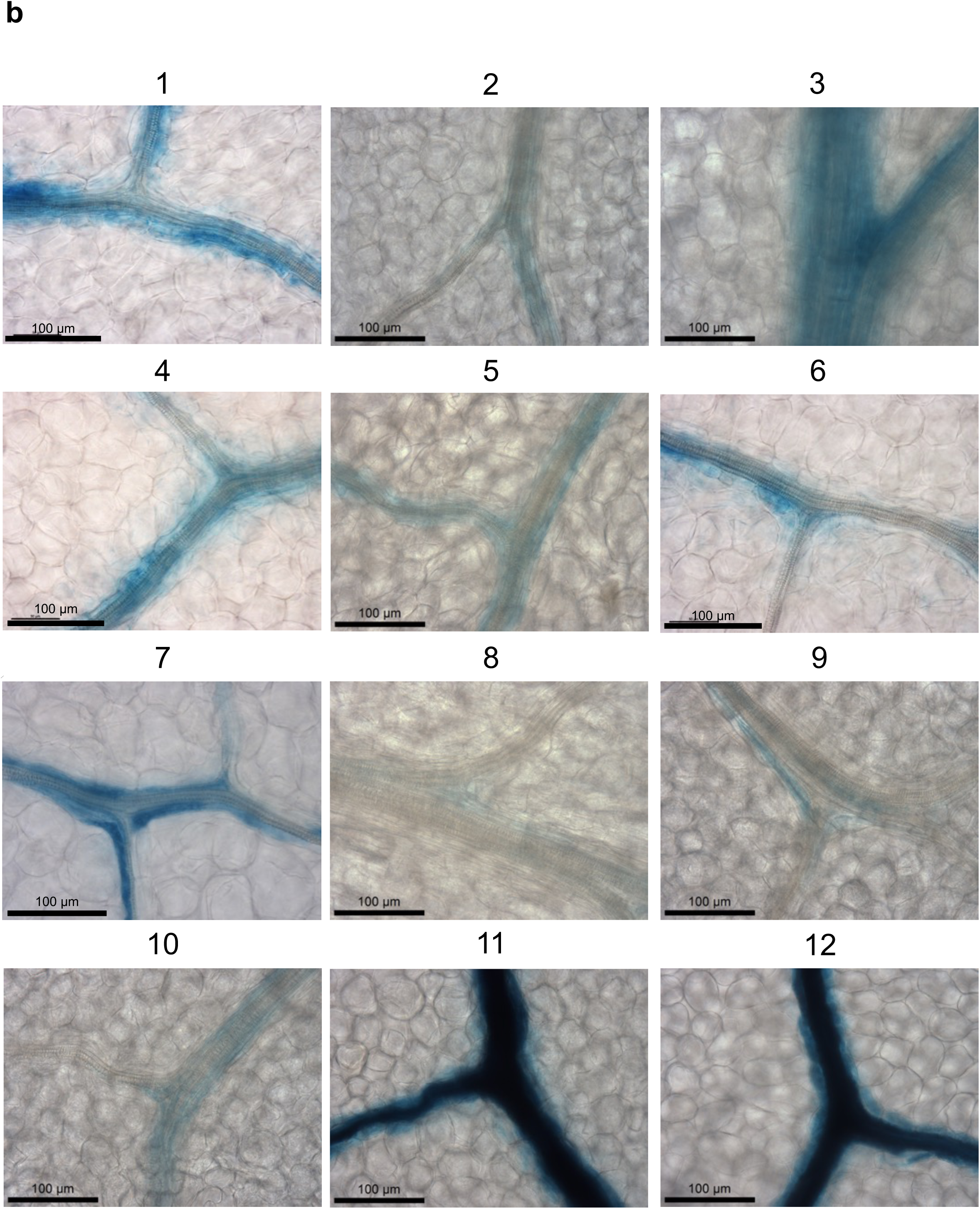
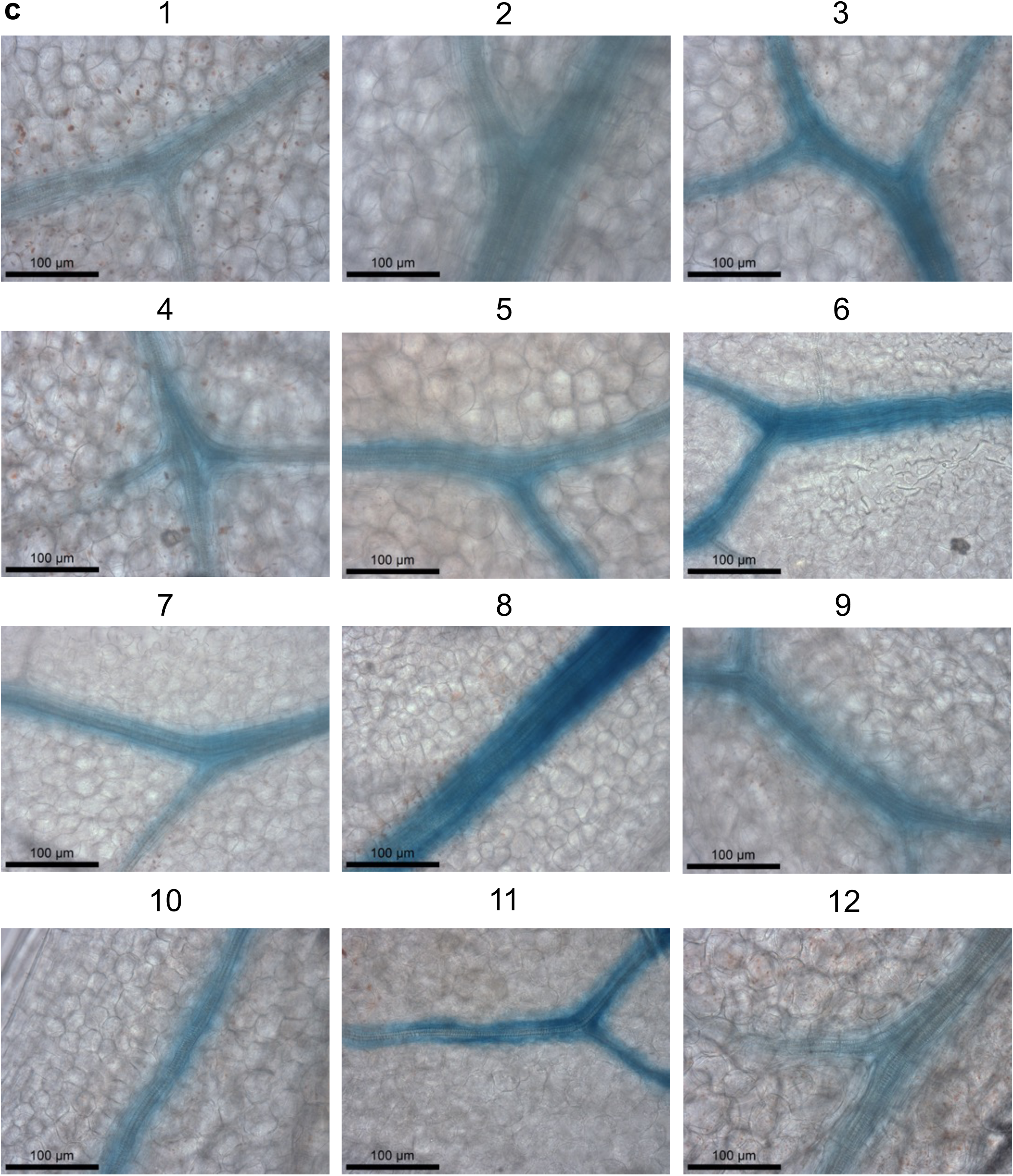
a) Deleting the MYB76 DHS leads to loss of GUS in the BS. Images from five independent transgenic lines, leaves were stained for 48 hrs. b) The *MYB76* DHS combined with the minimal 35SCaMV promoter generates preferential expression in the bundle sheath. Images from twelve independent transgenic lines. Leaves were stained for 72hrs. c) Oligomerizing the *MYB76* DHS combined with the minimal 35SCaMV promoter generates strong preferential expression in the bundle sheath. Images from twelve independent transgenic lines. Leaves were stained for 3hrs. Scale bars represent 100 µm.

**Supplementary Figure 8:**
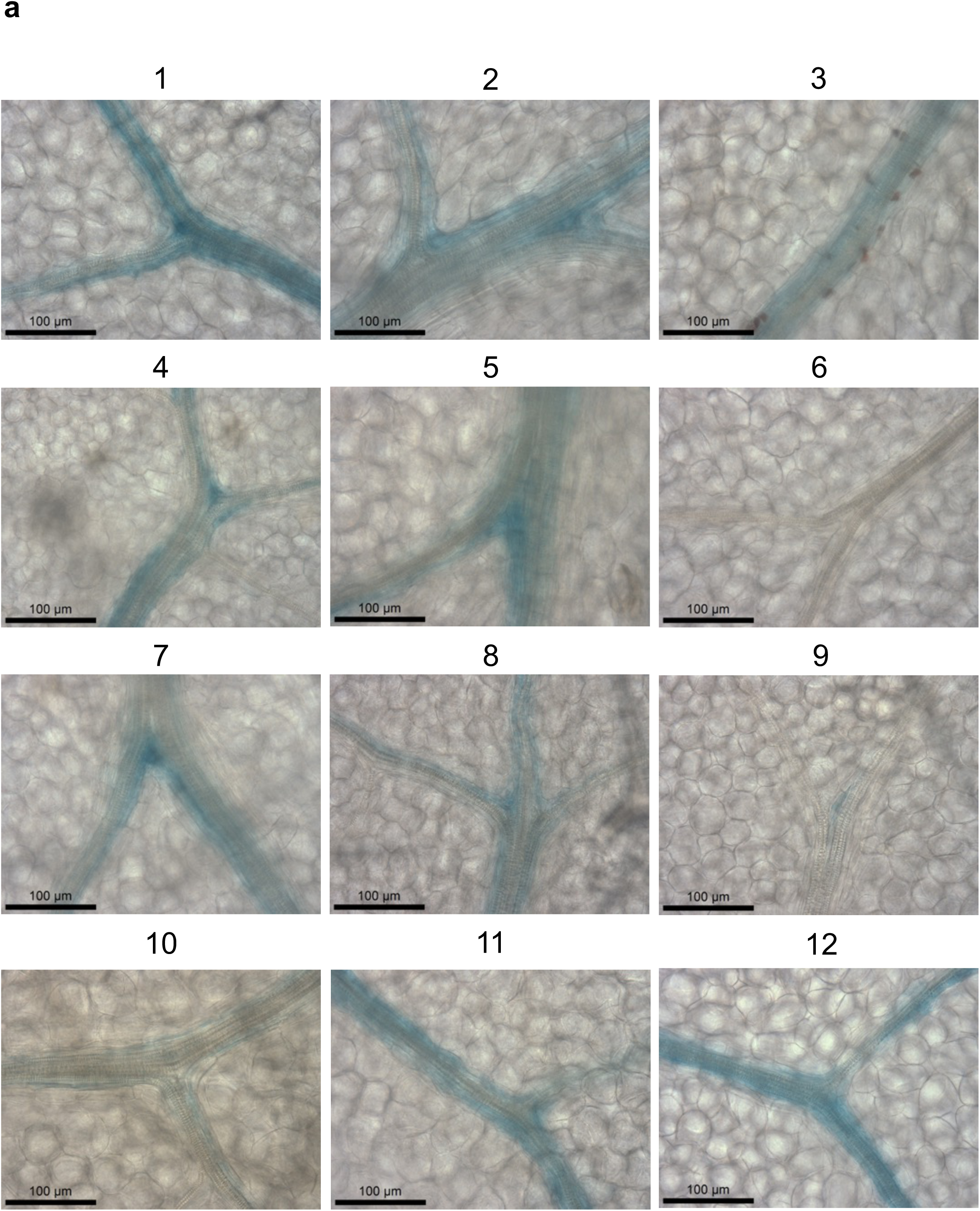
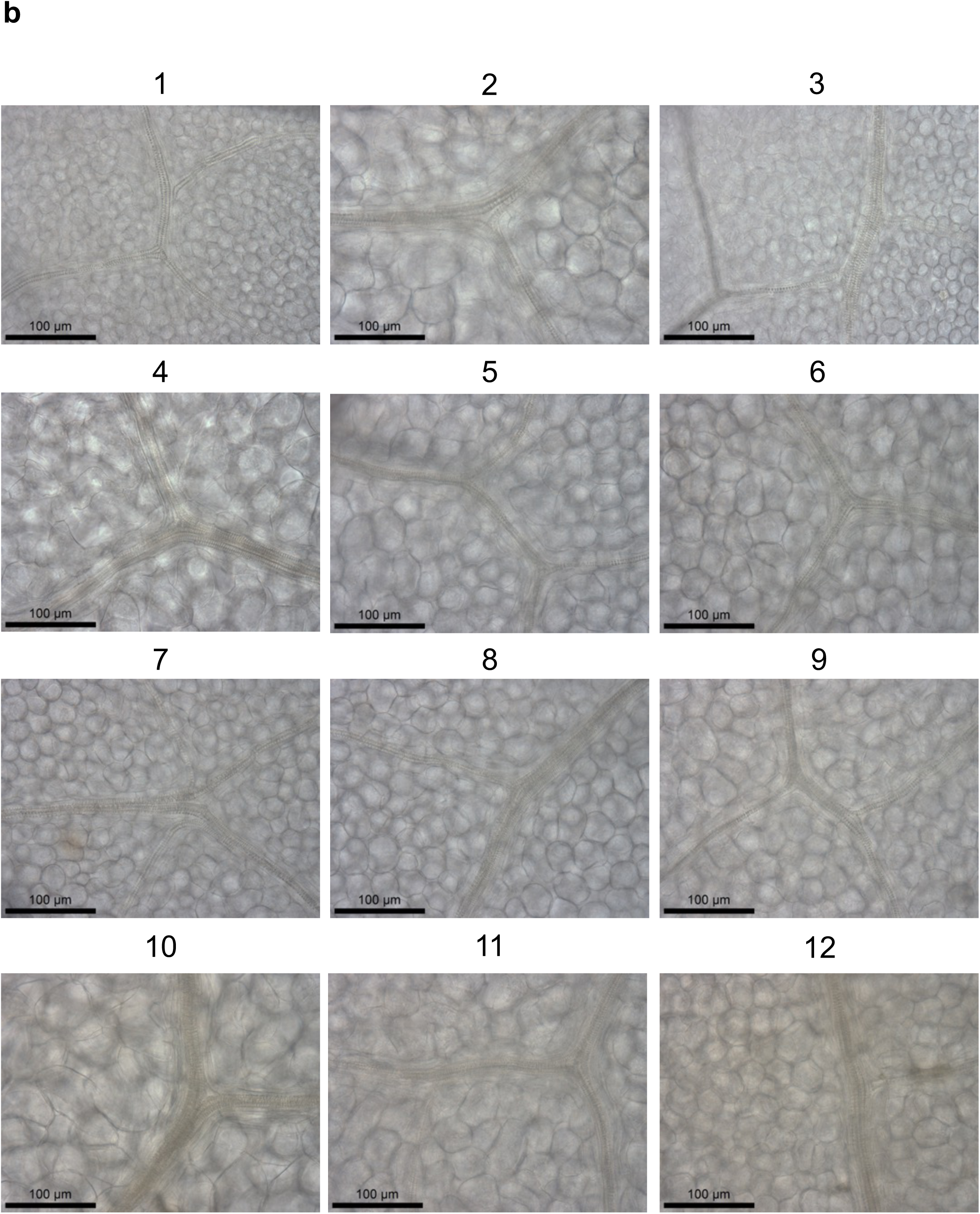
a) Mutation of motif 1 (TGGGCA) from the *MYB76* promoter does not abolish accumulation of GUS from the bundle sheath. Images from twelve independent transgenic lines. Leaves were stained for 48hrs. b) Mutation of motif 2 (TGCACCG) from the *MYB76* promoter motif leads to loss of GUS in the BS. Images from twelve independent transgenic lines. Leaves were stained for 48hrs. Scale bars represent 100 µm.

**Supplementary Figure 9:**
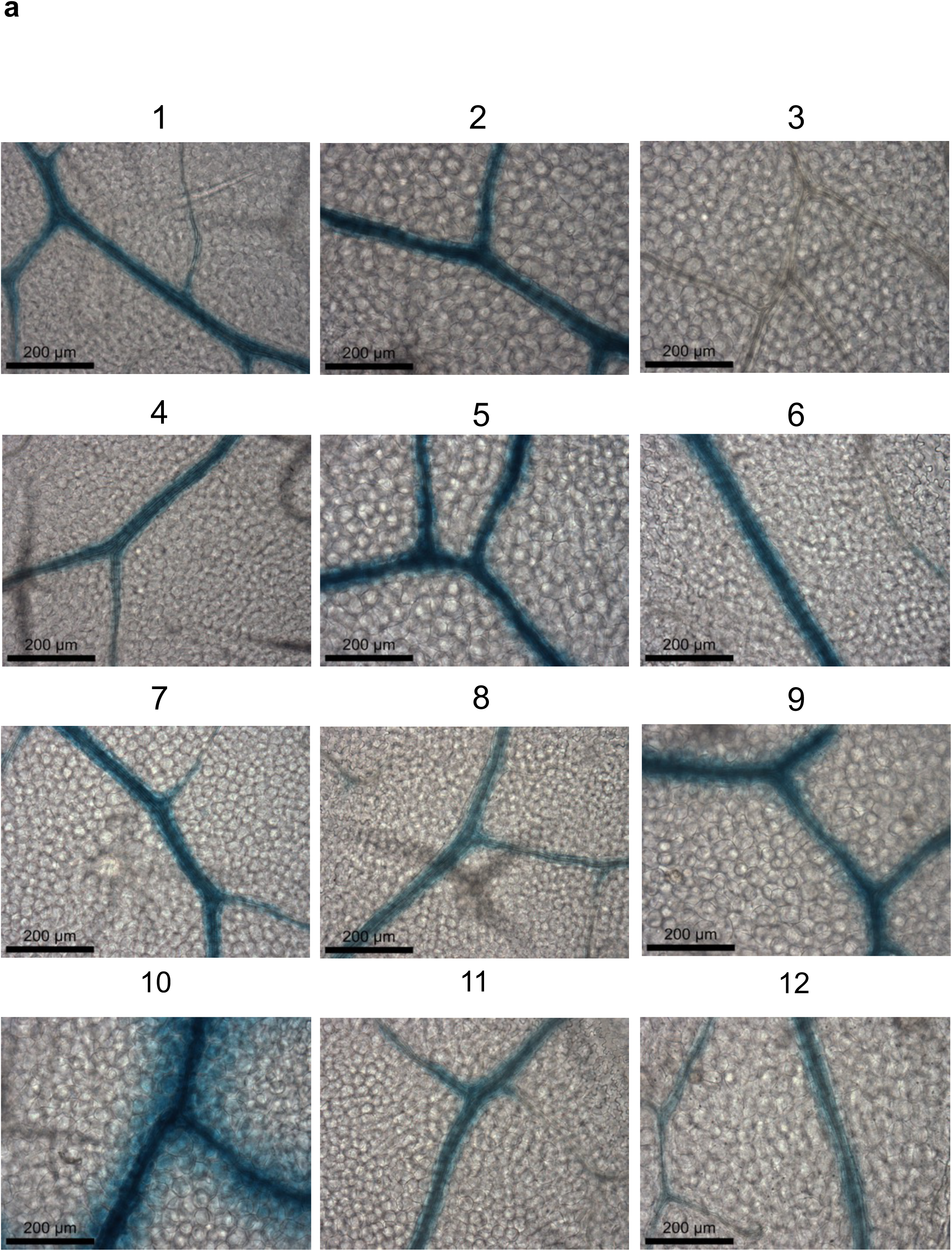
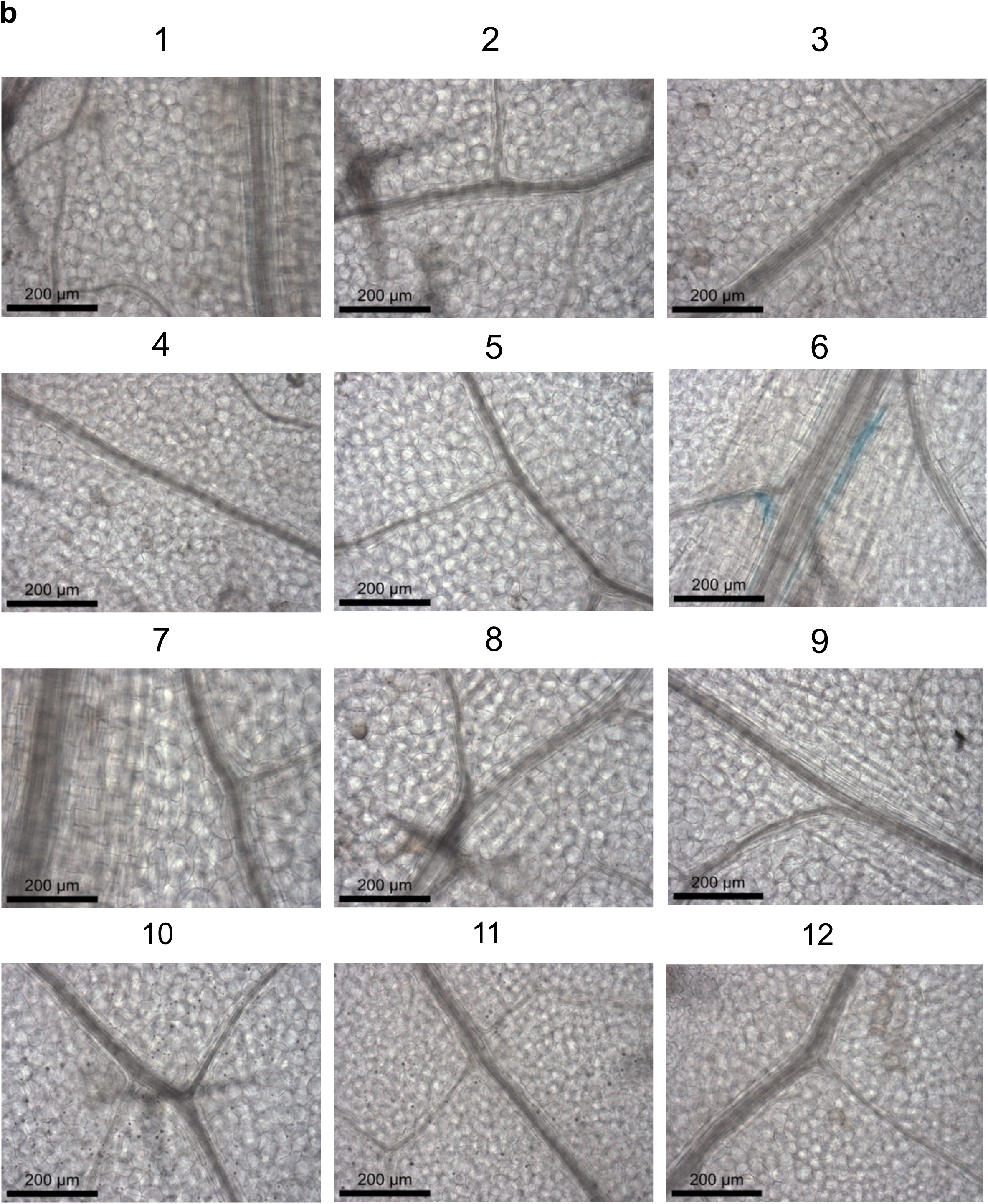
a) *MYB76* promoter and *gDNA* fused to GUS generated using Golden Gate cloning generates preferential expression in the bundle sheath. Images from twelve independent transgenic lines. b) Mutation of motif 2 (TGCACCG) in a full length *MYB76* promoter and gDNA translational fusion abolishes GUS accumulation. Images from twelve independent transgenic lines. Leaves were stained for 24 hrs. Scale bars represent 200 µm.

**Supplemental Figure 10:**
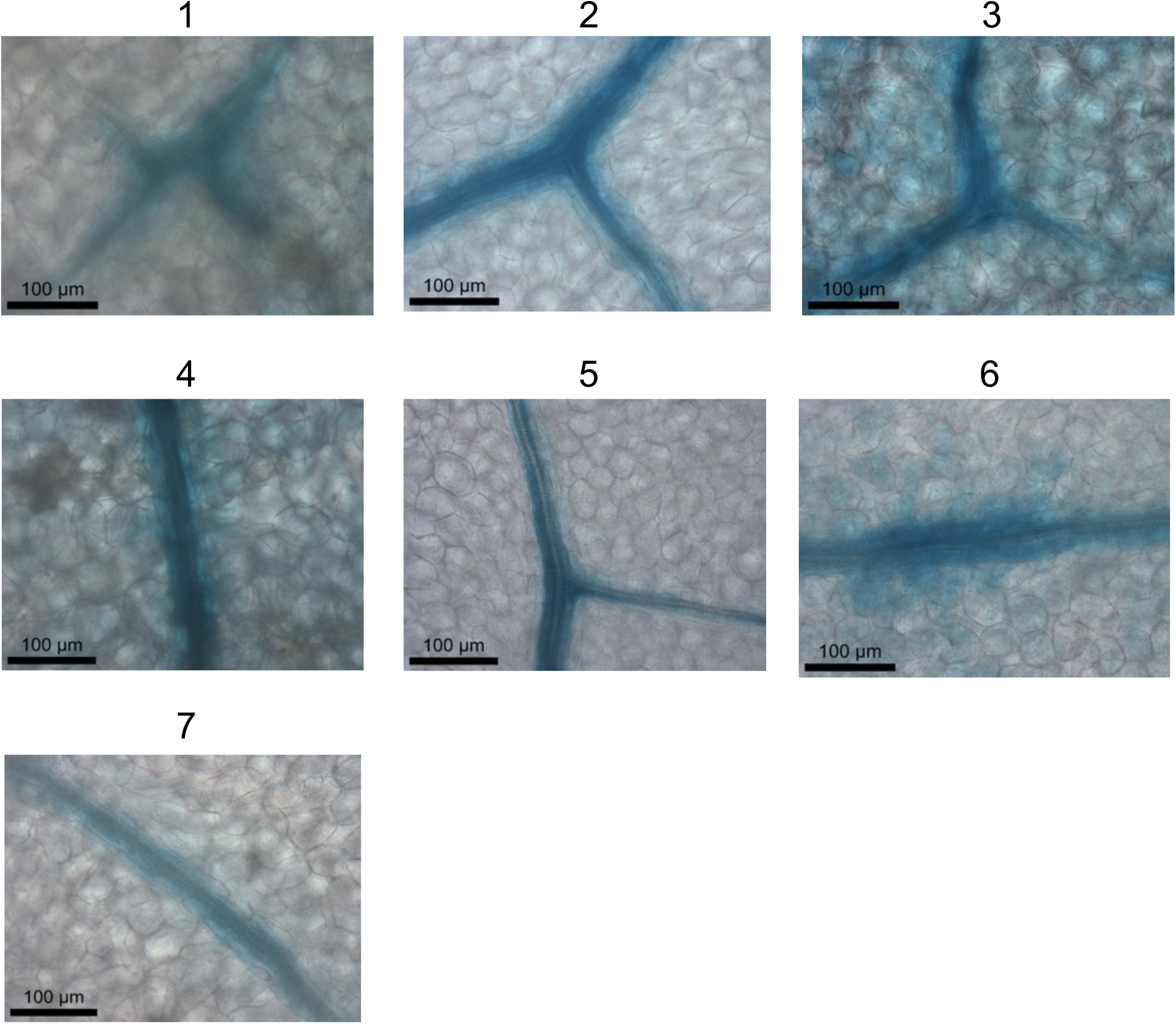
Two copies of the TGCACCG motif combined with ten upstream and ten downstream nucleotides within the context of the native *MYB76* promoter fused to the minimal 35SCaMV minimal promoter generate preferential expression in the bundle sheath. Images from seven independent transgenic lines. Leaves were stained for 86hrs. Scale bars represent 100 µm.

**Supplemental Figure 11.**
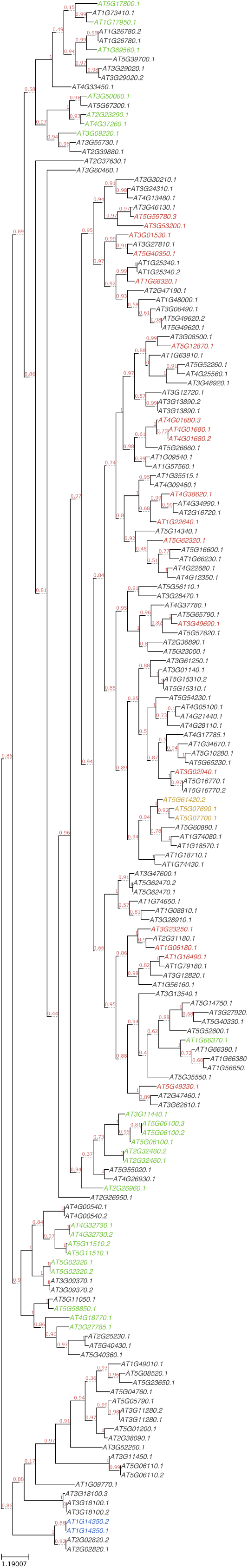
Phylogenetic tree of MYB transcription factors in *A. thaliana* based on amino acid sequence of whole proteins. Cluster 10 MYBs are coloured in green, cluster 18 MYBs are coloured in red and cluster 31 MYBs are coloured in blue. MYB transcription factors without defined binding motifs are in black and MYB28, MYB29 and MYB76 are coloured in gold.

**Supplemental Figure 12.**
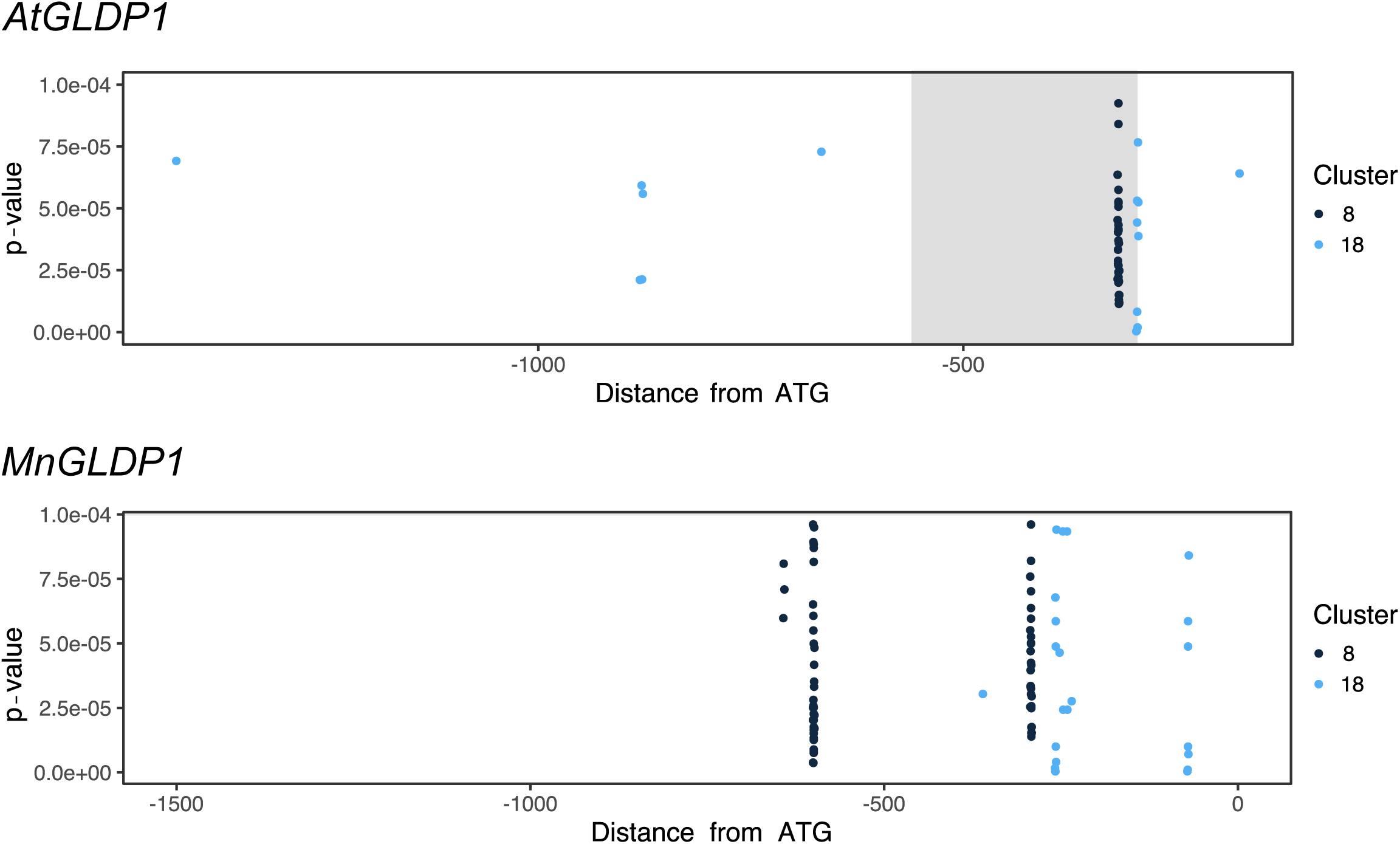
Cluster 8 and 18 transcription factor binding motifs within the promoters of *A. thaliana* and *M. nitens GLDP1* genes. The y axis shows p-values of matches between DHS sequence and motif PWMs and the x axis shows position of the motif centre relative to the translational start site. The grey box in the *AtGLDP1* promoter represents the V-box (Adwy et al., 2015).

## References

Adwy, W., Laxa, M. & Peterhansel, C. A simple mechanism for the establishment of C_2_- specific gene expression in Brassicaceae. Plant Journal 84, 1231–1238 (2015).

Adwy, W. et al. Loss of the M-box from the glycine decarboxylase P-subunit promoter in C_2_ Moricandia species. Plant Gene 18, 100176 (2019)

Agarwal, P.K. et al. Dehydration responsive element binding transcription factors and their applications for the engineering of stress tolerance. Journal of Experimental Botany 68, 2135–2148 (2017).

Akyildiz, M. et al. Evolution and function of a cis-regulatory module for mesophyll-specific gene expression in the C_4_ dicot Flaveria trinervia. The Plant Cell 19, 3391–3402 (2007).

Ali, S. & Taylor, W.C. The 3 ’ non-coding region of a C_4_ photosynthesis gene increases transgene expression when combined with heterologous promoters. Plant Molecular Biology 46, 325–333 (2001).

Aubry, S. et al. Transcript residency on ribosomes reveals a key role for the Arabidopsis thaliana bundle sheath in sulphur and glucosinolate metabolism. The Plant journal 78, 659– 673 (2014).

Bailey, T.L. et al. MEME Suite: tools for motif discovery and searching. Nucleic Acids Research 37, W202–W208 (2009).

Baima, S. et al. The expression of the Athb-8 homeobox gene is restricted to provascular cells in Arabidopsis thaliana. Development 121, 4171–4182 (1995).

Bhullar, S. et al. Strategies for Development of Functionally Equivalent Promoters with Minimum Sequence Homology for Transgene Expression in Plants: *cis*-Elements in a Novel DNA Context versus Domain Swapping. Plant Physiology 132, 988–998 (2003).

Bonke, M. et. al. APL regulates vascular tissue identity in Arabidopsis. Nature 426, 181–186 (2003).

Brown, N.J. et al. Independent and Parallel Recruitment of Preexisting Mechanisms Underlying C_4_ Photosynthesis. Science 331, 1436–1439 (2011).

Burow, M. et al. The Glucosinolate Biosynthetic Gene AOP2 Mediates Feed-back Regulation of Jasmonic Acid Signaling in Arabidopsis. Molecular Plant 8, 1201–1212 (2015).

Von Caemmerer, S., Quick, W.P. & Furbank, R.T. The development of C_4_ rice: Current progress and future challenges. Science 336 671–1672 (2012).

Castro-Mondragon J.A. et al. RSAT matrix-clustering: Dynamic exploration and re-dundancy reduction of transcription factor binding motif collections. Nucleic Acids Res 45, e119 (2017).

Clough, S.J. & Bent, A.F. Floral dip: a simplified method for Agrobacterium-mediated transformation of Arabidopsis thaliana. Plant Journal 16, 735–743 (1998).

Deplancke, B. et al. A Gene-Centered C. elegans Protein-DNA Interaction Network. Cell 125, 1193–1205 (2006).

Dey, N. et al. Synthetic promoters in planta. Planta 242, 1077–1094 (2015).

Du, Z. et al. agriGO: a GO analysis toolkit for the agricultural community. Nucleic Acids Res 38, W64–W70 (2010).

Egawa, C. et al. Differential regulation of transcript accumulation and alternative splicing of a *DREB2* homolog under abiotic stress conditions in common wheat. Genes & Genetic Systems 81, 77–91 (2006).

Engelmann, S. et al. The gene for the P-subunit of glycine decarboxylase from the C_4_ species *Flaveria trinervia*: analysis of transcriptional control in transgenic *Flaveria bidentis* (C_4_) and Arabidopsis (C_3_). Plant Physiology 146, 1773–1785 (2008).

Feike, D. et al. Characterizing standard genetic parts and establishing common principles for engineering legume and cereal roots. Plant Biotechnology Journal 17, 2234–2245 (2019).

Fernández-Calvo, P. et al. The Arabidopsis bHLH transcription factors MYC3 and MYC4 are targets of JAZ repressors and act additively with MYC2 in the activation of jasmonate responses. The Plant Cell 23, 701–715 (2011).

Fletcher, J.C. et al. Signaling of Cell Fate Decisions by CLAVATA3 in Arabidopsis Shoot Meristems. Science 19, 1911–1914 (1999).

Fornes, O., et al. JASPAR 2020: update of the open-access database of transcription factor binding profiles. Nucleic Acids Res., doi: 10.1093/nar/gkz1001 (2019).

Fryer, M.J. et al. Control of Ascorbate Peroxidase 2 expression by hydrogen peroxide and leaf water status during excess light stress reveals a functional organisation of Arabidopsis leaves. Plant Journal 33, 691–705 (2003).

Gallegos, J.E. & Rose, A.B. Intron DNA Sequences Can Be More Important Than the Proximal Promoter in Determining the Site of Transcript Initiation. The Plant Cell 29, 843– 853 (2017).

Gaudinier, A. et al. Enhanced Y1H assays for Arabidopsis. Nature methods 8, 1053–1055 (2011).

Gaudinier, A. et al. Identification of Protein–DNA Interactions Using Enhanced Yeast One-Hybrid Assays and a Semiautomated Approach. Methods Mol Biol 1610, 187–215 (2017).

Gigolashvili, T. et al. HAG2/MYB76 and HAG3/MYB29 exert a specific and coordinated control on the regulation of aliphatic glucosinolate biosynthesis in Arabidopsis thaliana. New Phytologist 177, 627–642 (2008).

Gowik, U. et al. *cis*-Regulatory elements for mesophyll-specific gene expression in the C_4_ plant *Flaveria trinervia*, the promoter of the C_4_ phospho *enol*pyruvate carboxylase gene. The Plant Cell 16, 1077–1090 (2004).

Grant, C.E., Bailey, T.L. & Noble, W.S. FIMO: scanning for occurrences of a given motif. Bioinformatics 27, 1017–1018 (2011).

Griffiths, H. et al. You’re so vein: bundle sheath physiology, phylogeny and evolution in C_3_ and C_4_ plants. Plant Cell Environ 36, 249–261 (2013).

Halkier, B.A. & Gershenzon, J. Biology and Biochemistry of Glucosinolates. Annual Review of Plant Biology 57, 303–333 (2006).

Heidstra, R., Welch, D. & Scheres, B. Mosaic analyses using marked activation and deletion clones dissect Arabidopsis SCARECROW action in asymmetric cell division. Genes and Development 18, 1964–1969 (2004).

Helariutta, Y. et al. The SHORT-ROOT Gene Controls Radial Patterning of the Arabidopsis Root through Radial Signaling. Cell 101, 555–567 (2000).

Hernandez-Garcia, C.M. & Finer, J.J. Identification and validation of promoters and *cis*-acting regulatory elements. Plant Science 217–218, 109–119 (2014).

Hesselberth, J.R. et al. Global mapping of protein-DNA interactions in vivo by digital genomic footprinting. Nature methods 6, 283–9 (2009).

Hibberd, J.M. & Covshoff, S. The Regulation of Gene Expression Required for C_4_ Photosynthesis. Annu. Rev. Plant Biol 61, 181–207 (2010).

Hibberd, J.M., Sheehy, J.E. & Langdale, J.A. Using C_4_ photosynthesis to increase the yield of rice - rationale and feasibility. Current Opinion in Plant Biology 11, 228–231 (2008).

Howe, G.A., Major, I.T. & Koo, A.J. Modularity in jasmonate signaling for multistress resilience. Annu. Rev. Plant Biol. 69, 387–415 (2018).

Imlau, A., Truernit, E. & Sauer, N. Cell-to-Cell and Long-Distance Trafficking of the Green Fluorescent Protein in the Phloem and Symplastic Unloading of the Protein into Sink Tissues. The Plant Cell 11, 309–322 (1999).

Jefferson, R.A., Kavanagh, T.A. & Bevan, M.W. GUS fusions: ß-Glucuronidase as a sensitive and versatile gene fusion marker in higher plants. EMBO Journal 6, 3901–3907 (1987).

Kajala, K. et al. Multiple Arabidopsis genes primed for direct recruitment into C_4_ photosynthesis. Plant Journal 69, 47–56 (2012).

Kinsman, E.A. & Pyke, K.A. Bundle sheath cells and cell-specific plastid development in Arabidopsis leaves. Development 125, 1815–1822 (1998).

Kirschner, S. et al. Expression of SULTR2;2, encoding a low-affinity sulphur transporter, in the Arabidopsis bundle sheath and vein cells is mediated by a positive regulator. Journal of Experimental Botany 69, 4897–4906 (2018).

Koroleva, O. et al. Patterns of solute in individual mesophyll, bundle sheath and epidermal cells of barley leaves induced to accumulate carbohydrate. New Phytol. 136, 97–104 (1997).

Leegood, R.C. Roles of the bundle sheath cells in leaves of C_3_ plants. Journal of Experimental Botany, 59, pp.1663–1673 (2008).

Levin, J.Z. & Meyerowitz, E.M. UFO - an Arabidopsis Gene Involved In Both Floral Meristem and Floral Organ Development. Plant Cell 7, 529–548 (1995).

Li, B. et al. Promoter-Based Integration in Plant Defense Regulation. Plant Physiology 166, 1803–1820 (2014).

Li, B. et al. Network-Guided Discovery of Extensive Epistasis between Transcription Factors Involved in Aliphatic Glucosinolate Biosynthesis. The Plant Cell 30, 178–195 (2018).

Liu, Q., et al. Two Transcription Factors, DREB1 and DREB2, with an EREBP/AP2 DNA Binding Domain Separate Two Cellular Signal Transduction Pathways in Drought- and Low-Temperature-Responsive Gene Expression, Respectively, in Arabidopsis. The Plant Cell 10, 1391–406. (1998)

Major, I.T. et al. Regulation of growth–defense balance by the JASMONATE ZIM-DOMAIN (JAZ)-MYC transcriptional module. New Phytologist 215, 1533–1547 (2017).

Malitsky, S. et al. The Transcript and Metabolite Networks Affected by the Two Clades of Arabidopsis Glucosinolate Biosynthesis Regulators. Plant Physiology 148, 2021–2049 (2008).

Mallmann, J. et al. The role of photorespiration during the evolution of C_4_ photosynthesis in the genus Flaveria. eLife 3, e02478 (2014).

Masucci, J.D. et al. The homeobox gene GLABRA2 is required for position-dependent cell differentiation in the root epidermis of Arabidopsis thaliana. Development 122, 1253–1260 (1996).

Matsukura, S. et al. Comprehensive analysis of rice DREB2-type genes that encode transcription factors involved in the expression of abiotic stress-responsive genes. Molecular Genetics and Genomics 283, 185–196 (2010).

Matsuoka, M. et al. The promoters of two carboxylases in a C_4_ plant (maize) direct cell-specific, light-regulated expression in a C_3_ plant (rice). The Plant Journal 6, 311–319 (1994).

Mustroph, A. et al. Profiling translatomes of discrete cell populations resolves altered cellular priorities during hypoxia in Arabidopsis. Proceedings of the National Academy of Sciences 106, 18843–18848 (2009).

Nakagawa, T. et al. Development of series of gateway binary vectors, pGWBs, for realizing efficient construction of fusion genes for plant transformation. J Biosci Bioeng 104, 34–41 (2007).

Nakamura, R.L. et al. Expression of an Arabidopsis Potassium Channel Gene in Guard Cells. Plant Physiology 109, 371–374 (1995).

Neph, S. et al. An expansive human regulatory lexicon encoded in transcription factor footprints. Nature 489, 83–90 (2012).

Otsuga, D. et al. REVOLUTA regulates meristem initiation at lateral positions. The Plant Journal 25, 223–236 (2001).

Patron, N.J. et al. Standards for plant synthetic biology: a common syntax for exchange of DNA parts. New Phytologist 208, 13–19 (2015).

Pruneda-Paz, J.L. et al. A Genome-Scale Resource for the Functional Characterization of Arabidopsis Transcription Factors. Cell Reports 8, 622–632 (2014).

Qin, F. et al. Regulation and functional analysis of ZmDREB2A in response to drought and heat stresses in Zea mays L. The Plant Journal 50, 54–69 (2007).

Quinlan, A.R. & Hall, I.M. BEDTools: a flexible suite of utilities for comparing genomic features. Bioinformatics 26, 841–842 (2010).

RStudio Team. RStudio: Integrated Development Environment for R http://www.rstudio.com/ (2015).

Reyna-Llorens, I. & Hibberd, J.M. Recruitment of pre-existing networks during the evolution of C_4_ photosynthesis. Philosophical Transactions of the Royal Society B: Biological Sciences 372, 2–7 (2017).

Reyna-Llorens, I. et al. Ancient duons may underpin spatial patterning of gene expression in C_4_ leaves. Proc. Natl Acad. Sci. USA 115, 1931–1936 (2018).

Robinson, J.T., et al. Integrative genomics viewer. Nature Biotechnology 29, 24 – 26 (2011).

Ruzicka, D.R. et al. The ancient subclasses of Arabidopsis ACTIN DEPOLYMERIZING FACTOR genes exhibit novel and differential expression. The Plant Journal 52, 460–472 (2007).

Sage, R. Environmental and evolutionary preconditions for the origin and diversification of the C_4_ photosynthetic syndrome. Plant Biology 3, 202–213 (2001).

Sakuma, Y. et al. Functional Analysis of an Arabidopsis Transcription Factor, DREB2A, Involved in Drought-Responsive Gene Expression. The Plant Cell 18, 1292–1309 (2006).

Salehin, M. et al. Auxin-sensitive Aux/IAA proteins mediate drought tolerance in Arabidopsis by regulating glucosinolate levels. Nature Communications 10, 4021 (2019).

Sarkar, A.K. et al. Conserved factors regulate signalling in Arabidopsis thaliana shoot and root stem cell organizers. Nature 446, 811–814 (2007).

Sarsby, J. et al. Mass spectrometry imaging of glucosinolates in Arabidopsis flowers and siliques. Phytochemistry 77, 110–118 (2012).

Schweizer, F. et al. Arabidopsis basic helix-loop-helix transcription factors MYC2, MYC3, and MYC4 regulate glucosinolate biosynthesis, insect performance, and feeding behavior. Plant Cell 25, 3117–3132 (2013).

Shatil-Cohen, A., Attia, Z. & Moshelion, M. Bundle-sheath cell regulation of xylem-mesophyll water transport via aquaporins under drought stress: a target of xylem-borne ABA? The Plant Journal 67, 72–80 (2011).

Sparks, E.E. et al. Establishment of expression in the SHORTROOT-SCARECROW transcriptional cascade through opposing activities of both activators and repressors. Developmental Cell 39, 585–596 (2016).

Sønderby I.E. et al. A systems biology approach identifies a R2R3MYB gene subfamily with distinct and overlapping functions in regulation of aliphatic glucosinolates. PLoS ONE 2, e1322 (2007).

Sønderby I.E. et al. A complex interplay of three R2R3 MYB transcription factors determines the profile of aliphatic glucosinolates in Arabidopsis. Plant Physiology 153, 348–363 (2010).

Takahashi, H. et al. The roles of three functional sulphate transporters involved in uptake and translocation of sulphate in Arabidopsis thaliana. Plant Journal 23, 171–182 (2000).

Theißen, G., Melzer, R. & Ruümpler, F. MADS-domain transcription factors and the floral quartet model of flower development: Linking plant development and evolution. Development 143, 3259–3271 (2016).

Thoma, S. et al. Tissue-specific expression of a gene encoding a cell wall-localized lipid transfer protein from Arabidopsis. Plant Physiology 105, 35–45 (1994).

Vainonen, J.P. et al. RCD1–DREB2A interaction in leaf senescence and stress responses in Arabidopsis *thaliana*. Biochemical Journal 442, 573–581 (2012).

Weber, E. et al. A Modular Cloning System for Standardized Assembly of Multigene Constructs. PLOS ONE 6, p.e16765 (2011).

Wickham, H. ggplot2: Elegant Graphics for Data Analysis. http://ggplot2.org/book/ (Springer, 2009).

Williams, B.P. et al. An untranslated *cis*-element regulates the accumulation of multiple C_4_ enzymes in *Gynandropsis gynandra* mesophyll cells. The Plant Cell 28, 454–465 (2016).

Williams, M. et al. Visualizing the distribution of elements within barley leaves by energy dispersive X-ray image maps (EDX maps). New Phytol. 125: 367–372 (2018).

Wiludda, C. et al. Regulation of the photorespiratory *GLDPA* gene in C_4_ *Flaveria*: an intricate interplay of transcriptional and posttranscriptional processes. Plant Cell 24, 137–151 (2012).

Wysocka-Diller, J.W. et al., Molecular analysis of SCARECROW function reveals a radial patterning mechanism common to root and shoot. Development 127, 595–603 (2000).

Yatusevich, R. et al. Genes of primary sulfate assimilation are part of the glucosinolate biosynthetic network in Arabidopsis thaliana. Plant Journal 62, 1–11 (2010).

Zhang, W. et al. Genome-Wide Identification of Regulatory DNA Elements and Protein-Binding Footprints Using Signatures of Open Chromatin in Arabidopsis. The Plant Cell 24, 2719–2731 (2012).

Zhang, T., Marand, A.P. & Jiang, J. PlantDHS: a databse for DNase1 hypersensitive sites in plants. Nucleic Acids Research 44, D1148–D1153 (2016).

